# The LINC complex component Kms1 and CENP-B protein Cbp1 cooperate to enforce faithful homology-directed DNA repair at the nuclear periphery in *S. pombe*

**DOI:** 10.1101/2025.04.12.648544

**Authors:** Alyssa Laffitte, Dongxu Lin, Yingzhen Jenny Tian, Na Liu, C. Patrick Lusk, Simon G. J. Mochrie, Megan C. King

## Abstract

While homologous recombination (HR) is often considered to be an error-free DNA repair mechanism, the fidelity of this pathway depends on the cell’s ability to engage the ideal template, the replicated sister chromatid. This is particularly challenging during repair of repetitive genome regions for which non-allelic sequences can errantly be used as templates. Here, we develop a model to study spontaneous DNA damage and repair that occurs at repetitive protein coding genes of the *S. pombe* flocculin family. We observe that genes encoding most members of this protein family constitutively reside at the nuclear periphery by virtue of their close proximity to binding sites for the CENP-B like protein, Cbp1. Tethering via Cbp1 to the nuclear periphery enhances the stability of the flocculin genes against intragenic recombination and restrains intergenic recombination between homoeologous repeat-encoding sequences. The LINC complex component Kms1 also antagonizes both intragenic and intergenic recombination at the flocculin genes as well as microhomology-mediated end-joining (MMEJ). Our observations suggest that *S. pombe* leverages nuclear compartmentalization to maintain the stability of repetitive genic regions at the nuclear periphery while association of DSBs with Kms1-containing LINC complexes enforces stringency to avoid mutagenic end-joining and use of the incorrect template during HR.

## Introduction

DNA double-stranded breaks (DSBs) are the most toxic type of DNA damage, leading to mutations, gross chromosomal rearrangements, and cell death if not efficiently and faithfully repaired (Cannan & Pederson, 2016). Cells use two major pathways to repair DSBs: canonical non-homologous end-joining (cNHEJ) and homologous recombination (HR). cNHEJ, the pathway of choice in the G1 phase of the cell cycle (Hustedt & Durocher, 2016), is promoted by the DSB end-binding Ku70/80 complex and the Lig4 DNA ligase, driving direct ligation of the broken DNA ends; cNHEJ frequently gives rise to small insertions and deletions (Sallmyr et al., 2024). In contrast, homologous recombination (HR) is up-regulated in the S and G2 phases of the cell cycle and represents a virtually error-free DSB repair pathway that engages the sister chromatid as a template for new DNA synthesis (Branzei & Foiani, 2008; Haber, 2018; Krejci et al., 2012; Li & Heyer, 2008; Lu et al., 2017). In order to achieve this, HR requires the resection of the 5’ ends of the DSB, creating 3’ single-stranded overhangs; these unstable ssDNA overhangs are protected by rapid recruitment of the abundant, ssDNA-binding Replication Protein A (RPA) complex (Symington, 2016). In yeasts, including the fission yeast *S. pombe*, further recruitment of Rad52 facilitates remodeling to load the recombinase Rad51, generating the nucleoprotein filament (New et al., 1998), which in turn catalyzes the search for the homologous sister chromatid (McIlwraith & West, 2008). Although challenging to monitor, the prevalent DSB repair mechanism in mitotically growing *S. pombe* cells, which reside primarily in the G2 phase of the cell cycle, is likely synthesis-dependent strand annealing (SDSA), a non-crossover form of HR (Elbakry & Löbrich, 2021; Haber, 2018; Helleday et al., 2007; Li & Heyer, 2008; Vines et al., 2022).

Although HR is considered the preferred pathway when the sister chromatid is available, its fidelity depends on the cell’s ability to identify the correct homologous sequence to use as a template. Following DNA synthesis, the sister chromatids are tethered together by cohesin complexes, limiting the need for an extensive homology search for productive, allelic HR; cohesin’s role in loop extrusion may also contribute to the homology search (Covo et al., 2010; Dumont et al., 2024; Piazza et al., 2021). However, faithful repair of tandem repeats nonetheless presents a challenging problem for the cell, including increasing the rate of replication fork stalling (Polleys et al., 2017; Triplett et al., 2024). During HR, strand invasion into a like repeat can drive intramolecular gene conversion or unequal sister chromatid exchanges if a cross-over occurs, driving repeat copy number changes (Hastings et al., 2009); SDSA (a non-crossover repair mechanism) can likewise lead to changes in repeat copy number (Elbakry & Löbrich, 2021; Wright et al., 2018). Use of an illegitimate repair templates at other genomic sites can be even more deleterious, leading to large deletions, insertions, loss-of-heterozygosity, or chromosomal translocations (Nian et al., 2024). However, for such repair outcomes to occur, the DSB must come into physical proximity with these non-allelic sequences (Renkawitz et al., 2014; Wang et al., 2017).

A key mechanism to protect repetitive regions from aberrant recombination involves nuclear compartmentalization. Fission yeast display the “Rabl configuration”, originally characterized by the centromeres and telomeres occupying opposite poles of the nucleus (Rabl, 1885) but further refined to include association of repeat-containing centromeres with the nuclear periphery adjacent to the spindle pole body (SPB) with repetitive telomeres tethered to the nuclear periphery distal to the SPB (Funabiki et al., 1993). In most biological contexts, the compartment at the nuclear periphery is enriched in both repetitive and silenced DNA, which are highly correlated (Padeken & Heun, 2014; Padeken et al., 2022); this relationship is often suggested to reflect heterochromatization of repetitive foreign DNA elements such as transposons including in fission yeast (Marsano & Dimitri, 2022; Murton et al., 2016; Padeken et al., 2022). However, actively expressed, multi-copy genes also reside at the nuclear periphery in yeasts, suggesting that these two facets can be uncoupled (Ahmed et al., 2010; Ahmed & Brickner, 2007; Brickner & Walter, 2004; Casolari et al., 2004; Chen & Gartenberg, 2014; Matsuda et al., 2017; Steglich et al., 2013; Sumner & Brickner, 2022). For example, Pol III transcribed genes such as tRNA genes and the 5S RNA genes (in *S. pombe*) cluster in 3D space in the nucleus and at the nuclear periphery in “TFIIIC bodies” (Cam et al., 2008; Chen & Gartenberg, 2014; Gallardo et al., 2019; Iwasaki et al., 2010; Lorenz et al., 2012). However, considering the nuclear periphery as a single compartment may be overly simplistic, as the nuclear envelope proteins Heh1/Lem2, Mps3/Sad1 and the nuclear pore complex (NPC) play distinct roles in orchestrating faithful DNA repair (Capella et al., 2021; Churikov et al., 2016; Conrad et al., 2008; Horigome et al., 2016; Horigome et al., 2014; Kalocsay et al., 2009; Kramarz et al., 2020; Moser et al., 2020; Nagai et al., 2008; Oza et al., 2009; Ryu et al., 2015)

Mechanistically, an emerging model is that pro-recombination factors are inhibited from loading onto DSBs within repeat-rich domains, with these lesions physically moving outside of the domain before HR proceeds; conceptually such a mechanism would help ensure use of the sister chromatid as a template for HR (Amaral et al., 2017; Anand et al., 2014; Caridi et al., 2017; Choi et al., 2022; Merigliano & Chiolo, 2021; Wang et al., 2017). For example, in budding yeast a DSB in the rDNA repeats fails to recruit Rad52 until the lesion moves outside of the nucleolus, permitting HR to proceed (Torres-Rosell et al., 2007); similar observations have been made at heterochromatic domains in *Drosophila* (Chiolo et al., 2011). In line with this model, the recruitment of HR factors is disfavored at DSBs associated with the nuclear lamina in mammalian cells (Lemaître et al., 2014) while DSBs in pericentric heterochromatin must move out of the heterochromatic domain in order to productively load Rad51 after replication in human cells (Tsouroula et al., 2016). Taken together, these studies suggest conserved mechanisms by which repetitive regions of the genome are subject to additional controls over DSB repair by HR.

Linker of Nucleoskeleton and Cytoskeleton (LINC) complexes and their constituent SUN and KASH domain proteins at the nuclear envelope appear also regulate DSB repair. We previously demonstrated that persistent DSBs associated with the SUN domain protein, Sad1, can engage with the KASH domain protein, Kms1, in the outer nuclear membrane to form a LINC complex that couples the DSB to dynamic cytoplasmic microtubules in *S. pombe* (Swartz, 2014). Moreover, LINC complexes have been found to influence DSB repair across eukaryotic models (Lawrence et al., 2016; Lottersberger et al., 2015; Oza et al., 2009; Shokrollahi et al., 2024). Interestingly, Kms1 also appeared to antagonize a poorly understood repair mechanism that allowed HR-deficient (*rad51Δ*) cells to survive (Swartz, 2014). One limitation of this study, like many, was that it primarily relied on the study of an inducible DSB that is irreparable in haploid cells rather than a naturally occurring DSBs that can be repaired using the sister chromatid as a template.

To overcome this challenge, here we develop a model to study spontaneous DNA damage and repair that occurs within a family of protein repeat-containing genes that encode *S. pombe* flocculins. We observe that the genes encoding most members of this protein family reside at the nuclear periphery, which we tie to nearby binding sites for the CENP-B like protein, Cbp1/Abp1. Tethering via Cbp1 to the nuclear periphery enhances the stability of the flocculin genes to both intragenic recombination within the protein repeat-encoding sequence and restrains intergenic recombination between homoeologous repeat-encoding sequences. In addition to the constitutive tethering of the flocculin genes to the nuclear periphery by Cbp1, the LINC complex component Kms1 appears to play a distinct role in enforcing genome stability. Using an assay to study alternative DNA repair mechanisms, we found that Kms1 restrains the use of microhomology-mediated end-joining (MMEJ), a mutagenic salvage DNA repair pathway. Taken together, our findings suggest a model in which Cbp1 and Kms1 both act to promote the stability of repetitive regions of the genome.

## Results

### *S. pombe* flocculin-like (*pfl*) genes are repetitive and localize to the nuclear periphery

To investigate the mechanisms that promote the stability of euchromatic, repetitive sequences, we sought to identify protein-coding genes containing internal repeats. We hypothesized that the *S. pombe* flocculin-like genes (*pfl*s), a rapidly evolving, expressed family of proteins (Fig. 1A and Supplemental Fig. 1) targeted to the cell surface that mediate cell-cell and cell-environment interactions (Kwon et al., 2012) could serve as such a model. Like the FLO genes of *S. cerevisiae* (also called adhesins) that are responsible for flocculation, biofilm formation and pseudohyphae (Verstrepen & Klis, 2006) the *pfl* genes encode similar protein structures that include a signal sequence, a variable number of highly related repeats of 35-36 amino acids (∼100 base pairs) and a ligand binding domain predominantly from one of two families – GLEYA and DIPSY (Linder & Gustafsson, 2008) (Fig. 1A, Supplemental Fig. 2). As the nuclear periphery has been broadly observed to stabilize repetitive regions of the genome, we first explored the subnuclear position of the *pfl* genes.

**Figure 1.**
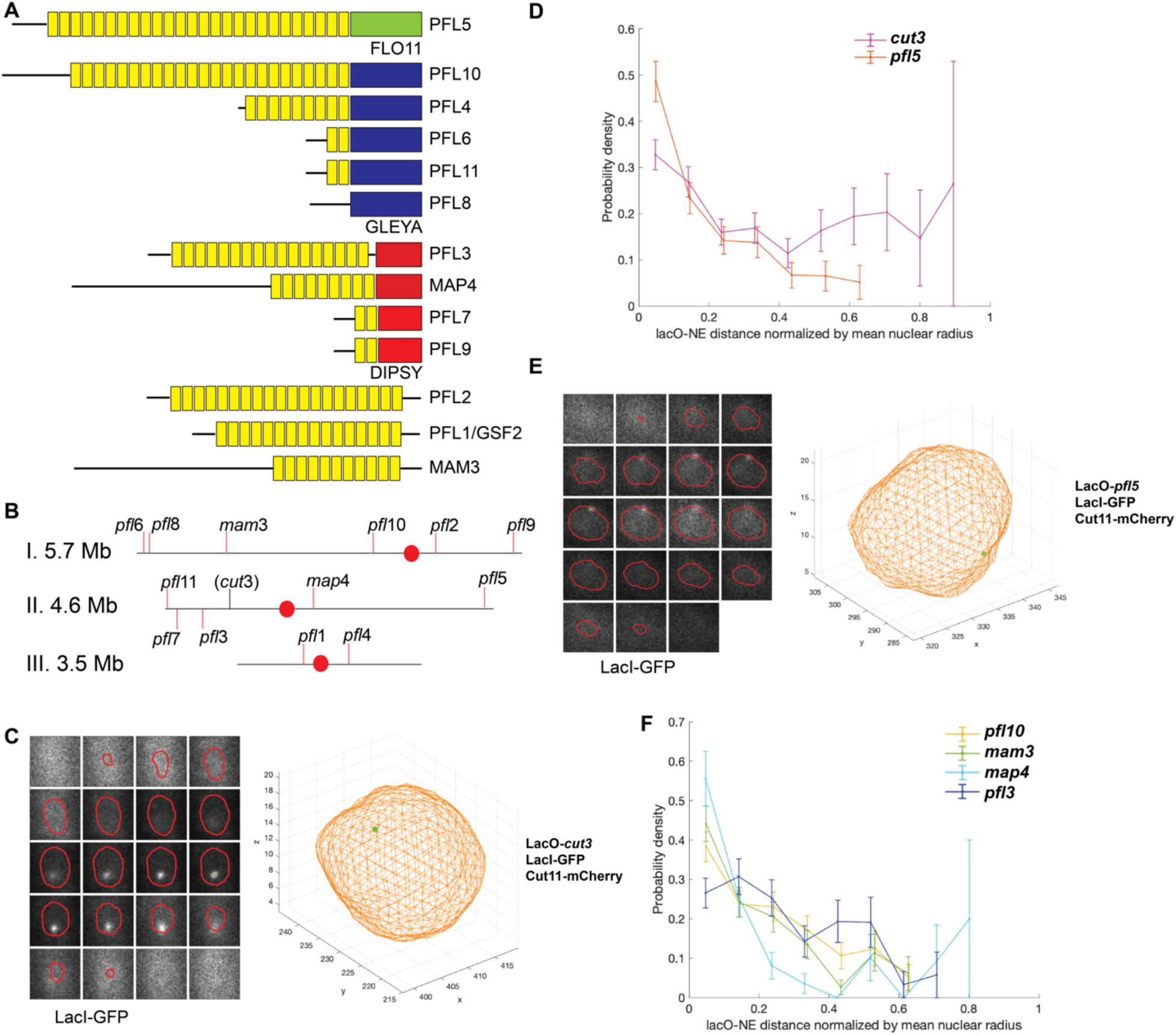
*S. pombe* flocculin-like (*pfl)* genes are a family of repetitive genes that localize to the nuclear periphery. A) *Pfl* genes encode similar structures that include a signal sequence (black line), a variable number of related repeats (yellow rectangles), and a ligand binding domain from one of two families: GLEYA (dark blue rectangles) and DIPSY (red rectangles). B) The position of the *pfl* genes in the *S. pombe* genome. Centromeres are represented by a red dot. C) The lacO/lacI-GFP system to visualize *pfl* genes within the nuclear volume defined by expressing a Cut11-mCherry fusion (Supplemental Fig. S3A). The lacO at *cut3* is centrally located within the nucleus. D) A probability density analysis shows that the lacO at *pfl5* is enriched at the nuclear periphery and depleted from the nuclear interior, while the lacO at *cut3* is broadly distributed across the nucleoplasm (n=224 nuclei for *plf5* and n=273 for *cut3*). E) A representative 3D reconstruction showing the lacO at *pfl5* at the nuclear periphery. F) A probability density analysis shows that the position of the lacO at each *pfl* gene tested (*pfl10* n=234*, mam3* n=189*, map4* n=101*, and pfl3* n=162) is biased towards the nuclear periphery.

To achieve this, we turned to the lacO/lacI-GFP chromatin tagging system to assess the location of *pfl* gene loci within the nuclear volume. We inserted a lacO array adjacent to *pfl* gene loci from each of the major families found across the chromosomes (Fig. 1A-B): *pfl5*, *pfl10*, and *pfl3* and the mating adhesin genes in the *pfl* family, *mam3* and *map4* (Mata & Bähler, 2006; Sharifmoghadam et al., 2006). As a comparison, we leveraged a euchromatic, non-*pfl* locus on that we characterized previously (*cut3*)(Zhao et al., 2016). To visualize the lacO array we expressed a tetramerization-deficient version of lacI fused to GFP and a nuclear localization signal, while we reconstruct the contour of the nuclear envelope based on the nucleoporin Cut11 expressed as an mCherry fusion, as in our prior work (Zhao et al., 2016)(Supplemental Fig. S3A). After acquiring z-stacks we employed custom MatLab image analysis pipelines we developed previously to fit the lacO/lacI-GFP signal to a Gaussian described by the point spread function and to reconstruct the nuclear surface (Zhao et al., 2016)(Supplemental Fig. S3A). In line with our prior observations, *cut3* was typically located within the nuclear interior, as illustrated by the representative reconstruction in Fig. 1C. To assess populations of cells, we plotted the probability distribution of distances from the center of the lacO/lacI-GFP Gaussian to the nearest point of the nuclear contour (normalized for nuclear size). Using this approach, we observed that *cut3* has a nearly uniform distribution throughout the nucleoplasm (Fig. 1E, pink). In comparison, *pfl*5 was found to be associated with the nuclear periphery as shown in the representative reconstruction in Fig. 1D and enriched at the nuclear periphery and depleted from the nuclear interior across a population of cells (Fig. 1E, brown). We observe a similar preference for a peripheral nuclear position for *map4* and *mam3* (Fig. 1F and Supplemental Fig. S3B-C). While *pfl10* and *pfl3* did not show the same extent of enrichment as *pfl5* at the nuclear periphery, they are nonetheless depleted from the interior of the nucleus compared to *cut3* (Fig. 1F and Supplemental Fig. S3D-E). Given their preferential association with the nuclear periphery, we employed the *pfl* gene family to further explore the relationship between nuclear position and the stability of repetitive protein-coding genes.

### Proteins involved in homologous recombination enforce stability of *pfl* genes

Given that direct repeats are prone to genetic instability due to intragenic recombination (Novarina et al., 2020; Rudin et al., 1989; Smith & Rothstein, 1999), we next asked if the *pfl* genes are prone to genetic instability. To assess this, we inserted a copy of the *ura4* gene within the direct repeats encoded by members of the *pfl* family (Fig. 2A). Cells were maintained on minimal medium without uracil before being subjected to fluctuation tests. To measure the stability of each *pfl* gene, cultures were inoculated into rich medium to allow for survival of cells lacking functional *ura4*. After saturation was achieved, the culture was plated onto media containing 5-Fluoorotic acid (5-FOA, a drug that produces a toxic metabolite in *ura4+* cells) to select for cells that had undergone spontaneous loss of *ura4* from the following *pfl* genes: *pfl3*, *pfl5*, *pfl10*, *mam3*, and *map4*. Counting the colonies that grew on 5-FOA relative to all cells plated (assessed from parallel plating onto rich medium) allowed us to measure the rate of loss of *ura4* (Supplemental Table S1). Given the range of direct repeat number inherent to the *pfl* gene family (Fig. 1A), we first asked if and how intrinsic instability is related to the number of repeats; a positive correlation was previously described between repeat copy number and intrinsic instability for the flocculin genes of *S. cerevisiae* (Verstrepen & Klis, 2006). As in this prior study, we observed a positive correlation between the frequency of 5-FOA-resistant colonies and repeat number (r^2^=0.8117), with *pfl5* (with 26 direct repeats) and *pfl10* (with 24 direct repeats) showing the highest rates of colony formation on 5-FOA (1.53 x 10^-5^ and 1.41 x 10^-5^, respectively) while *map4* and *mam3* with 9 and 10 direct repeats, respectively, showed lower rates of colony formation (1.59 x 10^-6^ and 5.25 x 10^-6^) (Fig. 2B). Importantly, the rate of spontaneous inactivating mutations in *ura4* at its endogenous locus are over three orders of magnitude less prevalent (<1 x 10^-8^)(Pai et al., 2023).

**Figure 2.**
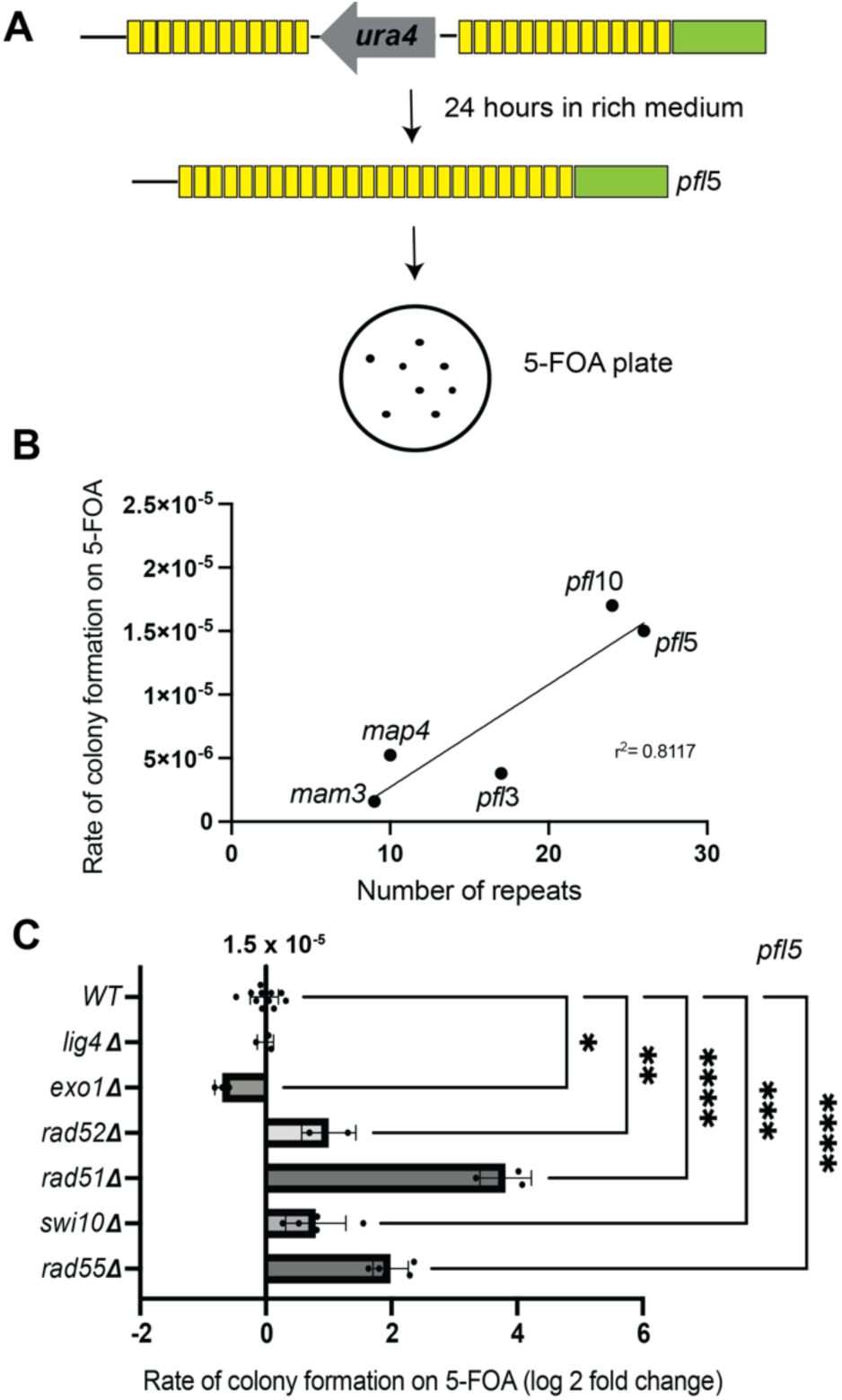
*Pfl* genes are genetically unstable. A) The stability of *pfl* genes can be monitored by the loss of *ura4* inserted into the direct repeats (represented by *pfl*5 here) and growth on selective plates containing 5-FOA. B) Across *pfl* loci, repeat number correlates with the rate of recombination (r^2^= 0.8117). C) HR-related DNA repair mechanisms impact the rate of *ura4* loss at *pfl*5. Loss of Lig4 shows no effect. Loss of Exo1 led to a decrease in recombination (p=0.0122 by one-way ANOVA with Dunnett correction for multiple comparisons). Loss of proteins that are involved in SSA or in the recombination step of HR, such as Rad52, Rad51, Rad55, and Swi10, cause a marked increase in the rate of recombination (p=0.0010, p<0.0001, p<0.0001, and p=0.0002, respectively). Data points are the natural log of the fold change in rate.

To provide insight into the mechanism(s) that drive the loss of *ura4* from within the *pfl* genes, we further investigated how disrupting a variety of established DNA repair factors influenced the rate of 5-FOA-resistant colony formation within *pfl5* (Fig. 2C). We examined genetic knockouts of Lig4 (the ligase driving canonical NHEJ), Exo1 (the exonuclease that drives long-range 3’ DSB resection upstream of homology-directed repair in fission yeast (Langerak et al., 2011; Leland et al., 2018)), the pro-HR factors Rad52 and Rad51, and Swi10 (the ERCC1 ortholog that promotes single-strand annealing between homologous DNA repeats (Zhang et al., 2014)). The *lig4Δ* strain behaved like WT, consistent with the preference of *S. pombe* to use HR during log phase growth (Ferreira & Cooper, 2004). Loss of Exo1 led to a ∼50% decrease in colony formation on 5-FOA, consistent with a role for long-range DSB resection upstream of homology-directed repair leading to loss of *ura4*. Somewhat surprisingly, loss of Rad52, which has roles in both loading of Rad51 onto resected DNA tails to drive HR as well in promoting single-strand annealing (SSA) of RPA-coated ssDNA (New et al., 1998), led to a nearly 3-fold increase in *ura4* instability. Loss of *rad51* itself led to an over 40-fold increase in rates of *ura4* loss with loss of *rad55* having a similar effect. It is important to note, however, that loss of RAD51 loading also leads to extensive degradation of resected ssDNA (Berti et al., 2020; Bonilla et al., 2020; Hashimoto et al., 2010); caution must therefore be exercised in concluding that this increase in rate reflects that homology-directed repair is dispensable for recombination-driven events observed in WT cells. To further explore this, we carried out diagnostic PCR on 5-FOA-resistant isolates from WT and *rad5111* cells. Compared to WT cells where we always see loss of *ura4* and some variability in repeat number (as expected) using PCR amplification across the *plf5* repeat region (16/16, Supplemental Fig. S4A-B), in most cases we fail to produce a product in *rad51Δ* cells (3/16, Supplemental Fig. S4C). There could be two explanations for this: 1) that the *ura4* gene is retained and therefore the product is too large to be amplified or 2) that the extensive degradation of resected DNA led to loss of the primer binding sites. Using a primer in the *ura4* cassette that can be amplified even in the absence of recombination, we observed a product in 2/13 isolates tested, suggesting that mutation of the *ura4* is a possible outcome in this background (Supplemental Fig. S4D). However, the most common outcome (11/16 isolates) is a regional loss of sequence around *pfl5* that includes loss of *ura4*. Thus, the increased rate of 5-FOA-resistant isolates in HR mutants likely stems primarily from extensive loss of sequence including *ura4*, while less common outcomes are successful reconstitution of *pfl5* repeats (possibly through single-strand annealing) or retention (and likely mutation) of *ura4*. Last, we observe that loss of Swi10 also drives an increase in colony formation on 5-FOA; given that its orthologs Rad10 (*S. cerevisiae*) and ERCC1 (human cells) have roles in flap cleavage to promote SSA (Osman et al., 2005), this suggests that either 1) despite *ura4* being embedded in direct repeats, this activity is not required, or 2) recombination promoted by Rad51 dominates in WT *S. pombe* in this context.

### A subset of *pfl* genes are tethered to the nuclear periphery via the CENP B-like protein, Cbp1

We next considered how the genomic context in which *pfl* genes reside might hint at the mechanism by which most *pfl* genes associate with the nuclear periphery. Like many rapidly evolving gene families, several of the *pfl* genes reside in large, multicopy segments in the subtelomeric regions of the genome (*pfl*5, *pfl*6, *pfl*7, *pfl*8, *pfl*9, *pfl*11; Fig. 3A); as telomeres are compartmentalized at the nuclear periphery in *S. pombe* (Matsuda et al., 2017), this could restrain their subnuclear position. While this could explain the observation that *pfl*5 localizes close to the nuclear periphery, the other *pfl* genes we characterized are located within the chromosome arms, including *pfl*3, *pfl*10, *map4*, and *mam3*, suggesting an additional mechanism at play. We were intrigued that the genes encoding *pfl*5 and *pfl*10 are directly adjacent to one of the thirteen full *Tf2* transposons in *S. pombe* (Bowen et al., 2003; Wood et al., 2002)(Fig. 3A). *Tf2* transposons are bound by the CENP-B homologs in fission yeast (Feng et al., 2012), which collectively recruit histone deacetylases to induce their repression (Lorenz et al., 2012); this activity is tied to physical clustering of *Tf2* transposons in “Tf bodies” (Cam et al., 2008; Feng et al., 2012). The CENP-B protein Cbp1/Abp1 also associates with abundant long terminal repeats (LTRs) lacking full transposon sequences and can silence nearby genes (Noma et al., 2004), regulate replication fork stability (Zaratiegui et al., 2011), and possibly more generally regulate regions of repetitive sequence (Daulny et al., 2016). We therefore leveraged existing data sets of Cbp1 binding peaks (Cam et al., 2008; Daulny et al., 2016) in the *S. pombe* genome to probe for a relationship with *pfl* genes. Using ChIP-on-chip data for Cbp1 (Cam et al., 2008) we found at least one Cbp1 binding peak within 600 bps of nearly all *Tf2*/LTRs as expected (264/282 or 93.6%) while a baseline prevalence for proximity within 600 bps of all protein-coding genes was 22.6% (1156/5123), likely reflecting the compactness of the *S. pombe* genome (Fig. 3B).

**Figure 3.**
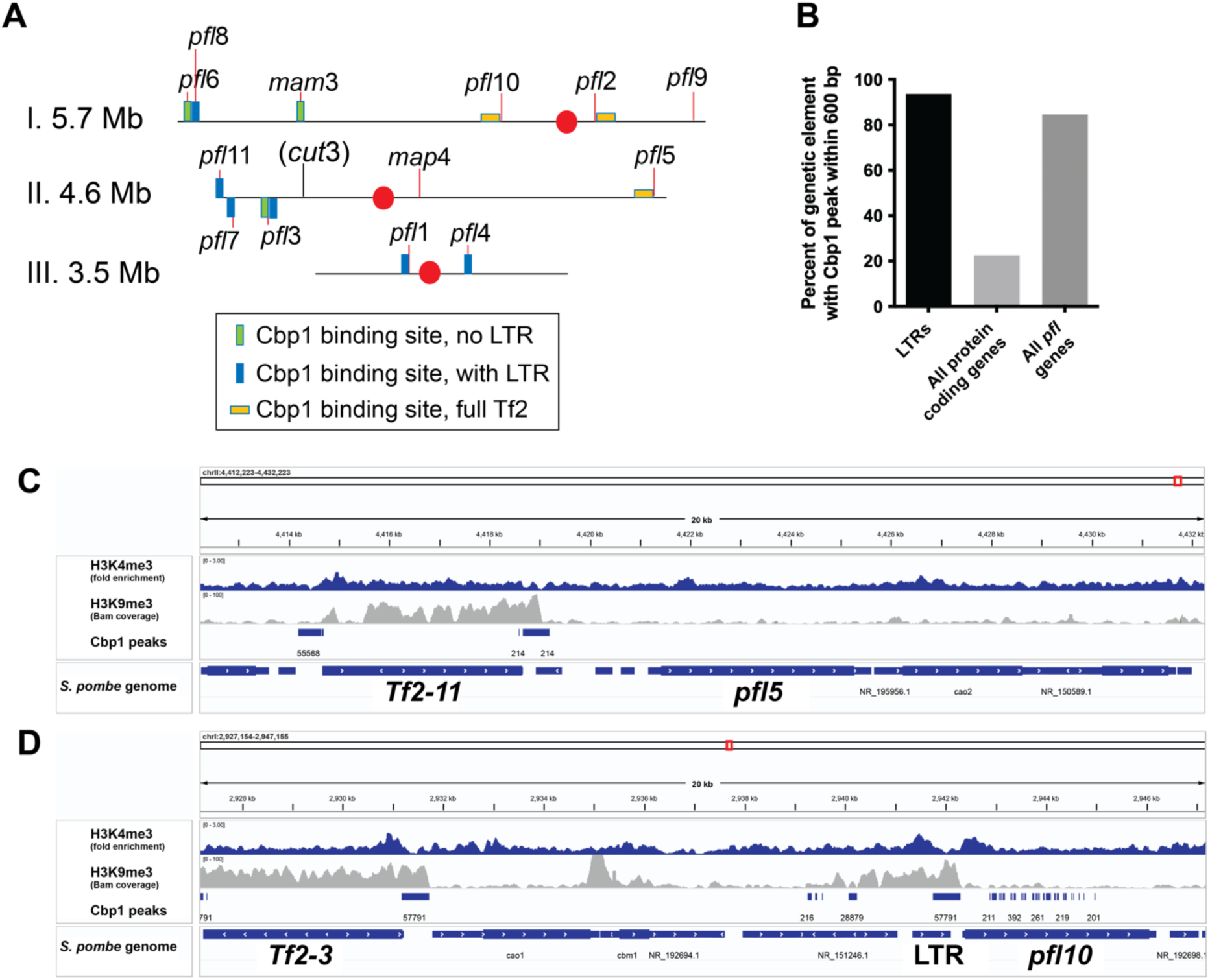
*Pfl* genes are nearby bindings sites for the CENP-B like protein, Cbp1. A) Most *pfl* genes are located nearby a Cbp1 binding site. Centromeres are represented by a red dot. B) 91% of LTR retrotransposons show a Cbp1 peak within 600bp, while only 20% of all protein coding genes show a Cbp1peak within 600bp. All *pfl* genes are more similar to LTRs (84.6%). C) Existing ChIP-Seq data (Daulny et al., 2016) shows Cbp1 peaks at *Tf2* transposons nearby *pfl5, and pfl10*. Data for histone marks H3K9me3 and H3K4me3 (DeGennaro et al., 2013; Shan et al., 2016) suggest the loci are actively transcribed.

Compared to this threshold, Cbp1 peaks were strongly over-represented near to *pfl* genes (within 600 bps for 11/13; 84.6%). Moreover, three *pfl* genes including *pfl2* (*Tf2-5*), *pfl5* (*Tf2-11*), and *plf10* (*Tf2-3*) are nearby full *Tf2* transposons (Fig. 3C,D). ChIP-seq data for Cbp1 confirms its association with these *Tf2* elements (Fig. 3C,D) (Daulny et al., 2016). The *pfl*10 locus also neighbors multiple additional Cbp1 binding peaks (Fig. 3D). Of note, despite their repetitive nature, these *pfl* genes show prevalent H3K4me3 (a mark of active transcription; (DeGennaro et al., 2013) and low levels of the repressive histone mark H3K9me3 (Shan et al., 2016)(Fig. 3C,D), consistent with their robust gene expression (Supplemental Fig. S1). This suggests that their proximity to Cbp1 binding sites and *Tf2* transposons is not sufficient to induce their silencing.

Given Cbp1’s ability to cluster *Tf2* transposons (Daulny et al., 2016; Murton et al., 2016; Shan et al., 2016), and the established relationship between transposable elements and the nuclear periphery (Marsano & Dimitri, 2022; Murton et al., 2016; Padeken et al., 2022), we next tested whether Cbp1 contributes to the tethering of *pfl* genes to the nuclear periphery. We compared the distribution of *pfl5*, *pfl10*, and *pfl3* (one representative of each family within the *pfl* genes) in WT and *cbp1Δ* genetic backgrounds using the lacO/lacI-GFP system (Fig. 4). We observed the clearest effect of the *cbp1Δ* condition on *pfl5*, where it was qualitatively obvious that in many cells the lacO at *pfl5* was more centrally positioned in the nucleus in the *cbp1Δ* condition compared to the WT control (Fig. 4A) as reinforced by representative reconstructions (Fig. 4B). The probability distribution for the lacO at *pfl5* is also strongly influenced by deletion of Cbp1, with a loss of enrichment at the nuclear periphery (Fig. 4C). Of note, while *cbp111* cells tend to have slightly larger nuclear volumes (on average an increase of 16% compared to WT cells (Supplemental Fig. S5A)), this can account for a shift of less than a single bin of the probability distribution analysis and therefore cannot account for the profound shift in *pfl5* position. While less dramatic for *pfl3* (Fig. 4D) and *pfl10* (Fig. 4E), both of these loci also became more randomly distributed throughout the nucleoplasm in cells lacking Cbp1. These results suggest that Cbp1 contributes to the peripheral tethering of at least a subset of the *pfl* genes.

**Figure 4.**
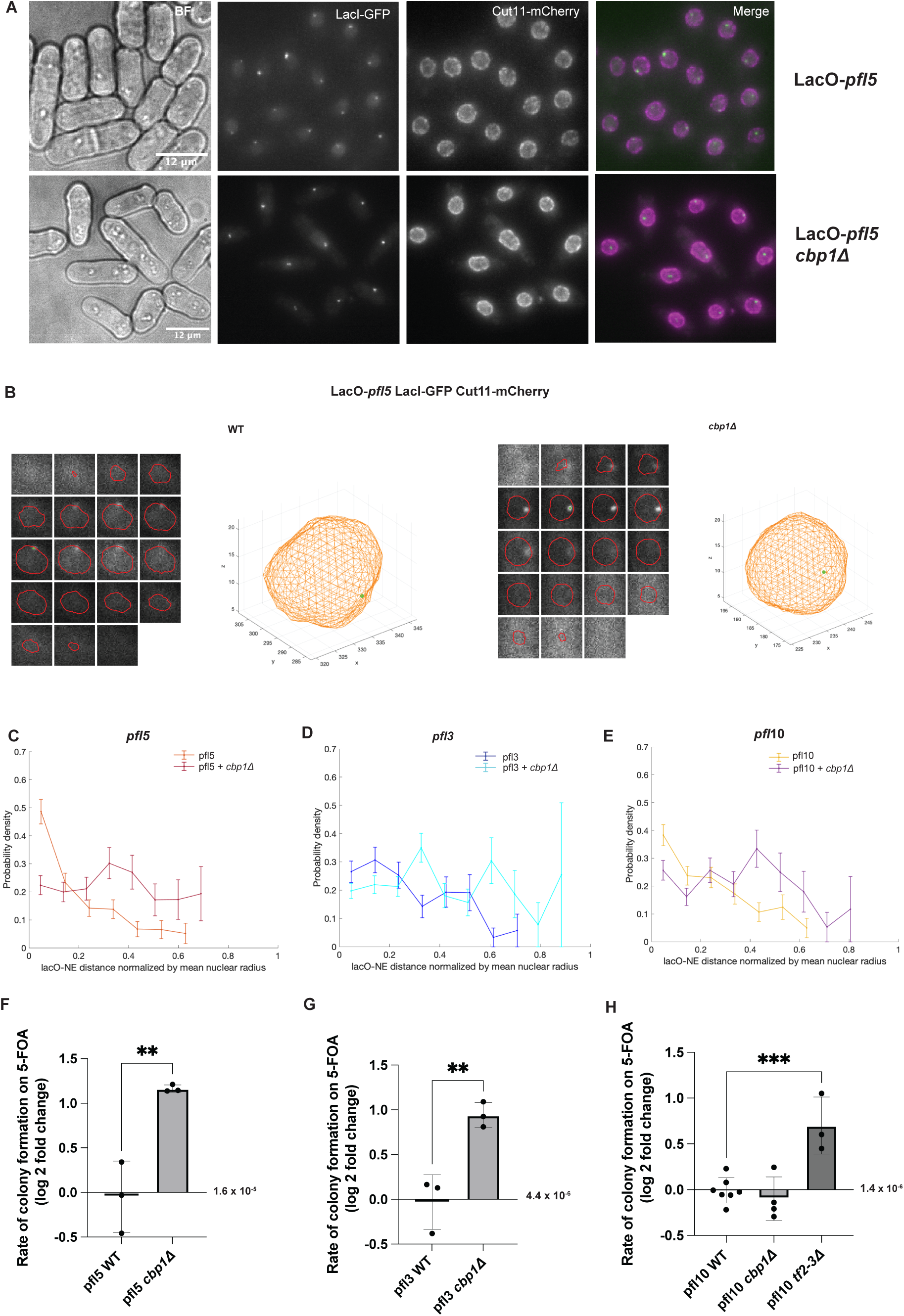
A subset of *pfl* genes are tethered to the nuclear periphery via the CENP B-like protein, Cbp1. A-B) Using the lacO/lacI-GFP system, we observe that loss of *cbp1* releases *pfl5* from the nuclear periphery. C-E) Probability density functions of the distance from the lacO to the nuclear envelope for *pfl5, pfl3* and *pfl10* with and without Cbp1. Data for WT cells replotted from Fig. 1. *plf5* WT n=224, *pfl5 cbp1*Λ n= 172, *plf3* WT n=162, *plf3 cbp1*Λ n=235, *pfl10* WT n=234, *pfl10 cbp1*Λ n=181. F-G) Loss of *cbp1* causes an increase in the rate of recombination at the *pfl5* and *pfl3* loci. A pairwise t-test indicates p=0.0075 for *pfl5* and p=0.0064 for *pfl3.* H) Loss of *cbp1* does not impact the rate of recombination at *pfl*10. However, deletion of *Tf2-3* nearby the *pfl*10 locus leads to an increase in recombination. A one-way ANOVA with Dunnett’s multiple comparison test indicates p=0.0007. Data points are the natural log of the fold change in rate.

### Cbp1 maintains the stability of *pfl* genes

Cbp1 was previously demonstrated to restrain the recombination of a *Tf1* transgene into remnant *Tf1* LTRs in the *S. pombe* genome (Feng et al., 2012). This finding, in addition to our observation that Cbp1 contributes to the tethering of *pfl* genes to the nuclear periphery, led us to ask if Cbp1 also influences the stability of the *pfl* genes. We again leveraged the intragenic recombination assay. Genetic deletion of Cbp1 increased the rate of recombination at both *pfl*5 and *pfl*3 (Figs. 4F-G and Supplemental Table S1); for *pfl*5 the rate of recombination increased from 1.53 x 10^-5^ in the WT cells to 5.06 x 10^-5^ in the *cbp1Δ* cells while for *pfl3* the rate of recombination increased from 4.39 x 10^-6^ in WT cells to 1.14 x 10^-5^ in the *cbp1Δ* cells. By contrast, we found that genetic deletion of Cbp1 did not affect genomic stability at *pfl10* (Fig. 4, Supplemental Table S1). Consistent with a specific effect of Cbp1 on instability of *plf5* and *pfl3* but not *pfl10*, we failed to observe an increase in spontaneous DNA damage globally as assessed by monitoring the number of Rad52-GFP foci in *cbp1Δ* cells (Supplemental Fig. S5B). To examine if Cbp1 might act indirectly to drive instability of the *pfl* genes by influencing their expression we carried out RT-qPCR. We observe a subtle up-regulation of *pfl5* in *cbp1Δ* cells (< 3-fold) while there was little effect on *pfl3* and *pfl10.* To directly investigate if *pfl5* expression levels can influence its stability we integrated the heterologous, regulatable *nmt41* promoter that drives moderate expression in rich media conditions and high expression in minimal media lacking thiamine (Iwaki & Takegawa, 2004) upstream of the *pfl5* coding sequence. However, we observed that induction led to a subtle decrease in instability (Supplemental Fig. S5C). While *pfl10* stability was not affected by loss of Cbp1, we found that deleting the nearby binding site for CENP-B proteins (an intact *Tf2* transposon, *Tf2-3*) was sufficient to drive a two-fold increase in the rate of recovering 5-FOA-resistant isolates (Fig. 4H and Supplemental Table S1). Taken together, these observations support the hypothesis that tethering to the nuclear periphery through Cbp1 for *pfl5* and *pfl3*, and possibly in concert with other CENP-B proteins for *pfl10*, contributes to the stability of *pfl* genes independent of heterochromatin state and/or their expression level.

### Cbp1 and the LINC complex cooperate to enforce genetic stability of the *pfl* genes

We previously reported that the Sad1-Kms1 LINC complex couples persistent DSBs at the nuclear periphery to dynamic cytoplasmic microtubules (Swartz, 2014). We therefore interrogated if loss of *kms1* impacts the stability of *ura4* embedded in the direct repeats of the constitutively peripheral *pfl5* gene. We found that loss of Kms1 increases the rate of colony formation on 5-FOA from 1.53 x 10^-5^ for WT to 2.39 x 10^-5^ for *kms111* cells (Fig. 5A and Supplemental Table S1). While this increase did not reach statistical significance using an ANOVA test with a Tukey correction for multiple comparisons when comparing all conditions (p=0.15), we obtain a value of p=0.02 using a pairwise t-test of these two conditions alone. To ask whether Kms1 acts in the same pathway as Cbp1, we compared the effect of the single and combined *kms111* and *cbp111* alleles. Similar to the results in Fig. 4F, loss of Cbp1 alone leads to an increase of more than 2-fold in the rate of acquiring 5-FOA resistance (the rate for *cbp111* was 5.06 x 10^-5^; Fig. 5A and Supplemental Table S1). There seems to be no additive effect between the two alleles, as *kms111 cbp111* cells show a rate of 4.73 x 10^-5^), suggesting that these two factors likely function in the same pathway. Thus, Kms1 likely cannot influence the intragenic repair outcome when *pfl5* is released from the nuclear periphery in *cbp111* cells.

**Figure 5.**
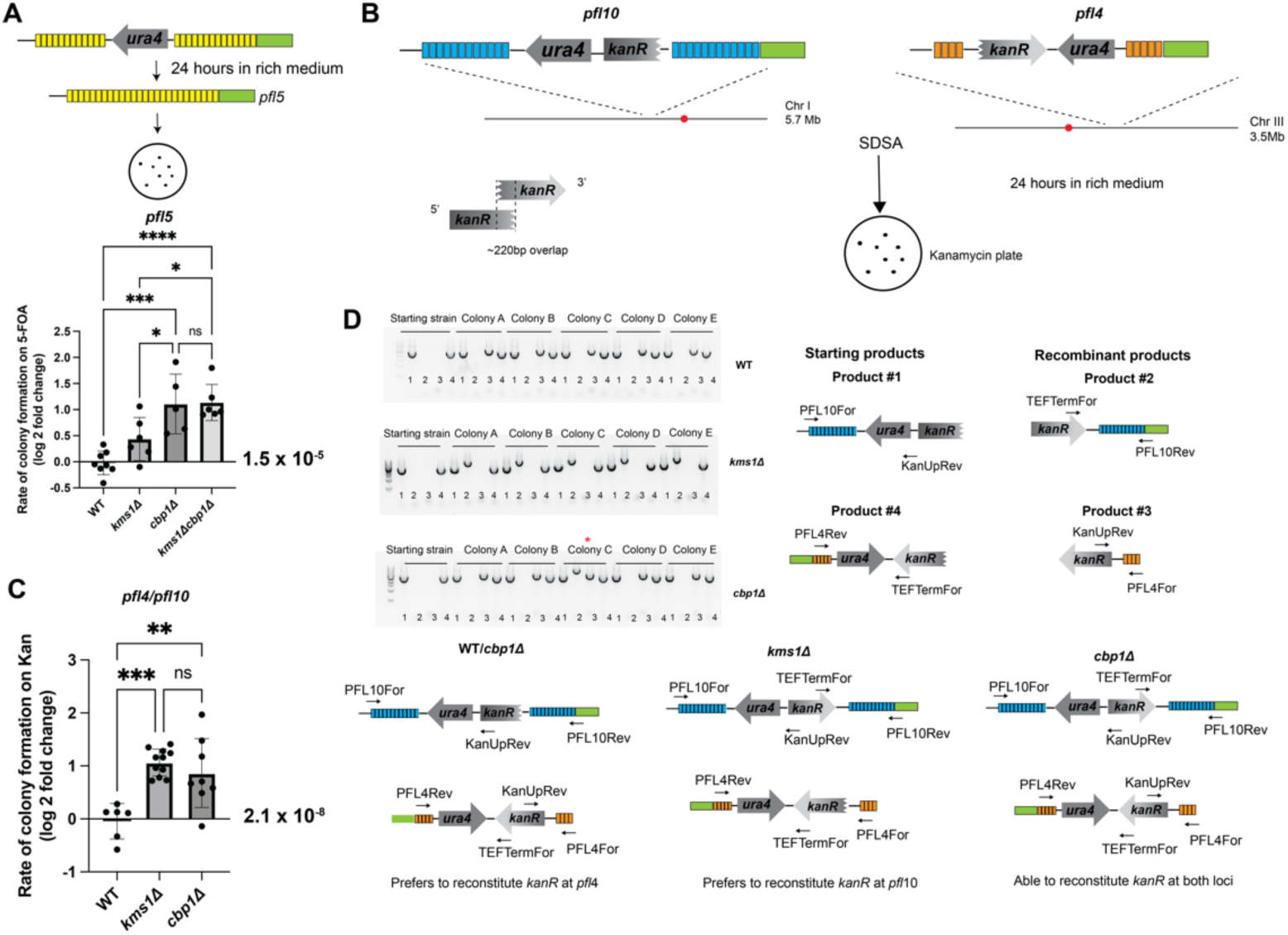
Cbp1 and the LINC complex cooperate to enforce genetic stability of the *pfl* genes. A) Loss of *kms1* increases the rate of colony formation on 5-FOA in the intragenic recombination assay at *pfl5*, although to a lesser extent than loss of *cbp1* (p=0.0002 by one-way ANOVA with a Tukey correction for multiple comparisons). Data points are the natural log of the fold change in rate. B) An assay to measure recombination between two homoeologous regions. Potential SDSA products are drawn out, along with PCR primers to amplify across the loci to determine the mechanism of acquiring Kan resistance. C) Loss of Cbp1 or Kms1 increases recombination between homoeologous loci. The rate of colony growth on Kan plates increases in *cbp1Δ* or *kms1Δ* cells, suggesting that both genes enforce stringency in choosing the correct template during repair by homologous recombination. (p<0.0001 for *kms1Δ* and p=0.0016 for *cbp1Δ* by one-way ANOVA with Dunnett’s correction for multiple comparisons.) Data points are the natural log of the fold change in rate. D) PCR products between the *kanR* coding sequence and locus specific primers provided insight into how the cells reconstituted *kanR*. WT and *cbp1Δ* cells prefer to gain the 5’ side of *kanR* at *pfl*4 (“recombinant product” #3), suggesting *kanR* was reconstituted at *pfl*4. However, *kms1Δ* colonies gained the 3’ side of *kanR* at *pfl*10 (“recombinant product” #2), suggesting *kanR* was reconstituted at *pfl*10. *cbp1Δ* colonies are also capable of reconstituting *kanR* at both loci (Colony C, see asterisk).

The LINC complex also promotes high fidelity homologous chromosome pairing during meiosis across model systems (Conrad et al., 2008; Horn et al., 2013; Lee et al., 2015; Sato et al., 2009; Varas et al., 2015) possibly by transmitting cytoskeletal forces to the chromosomes during meiotic prophase and dissolving non-homologous chromosome pairs (Sato et al., 2009). We therefore considered the possibility that the LINC complex could enforce the use of the homologous repair template (the sister chromatid) during DNA repair by homologous recombination. To test this hypothesis, we created an assay to measure illegitimate recombination between two “homoeologous” (similar, but not identical) loci (Fig. 5B). We inserted overlapping halves of the Kanamycin-resistance gene cassette (*kanR*) into two homoeologous loci from the same *pfl* gene subfamily with the highest extent of identity (Supplemental Fig. S6) that reside on separate chromosomes, *pfl10* (chromosome I) and *pfl4* (chromosome III); the 5’ and 3’ regions of *kanR* contained a ∼220bp overlap; the *ura4* marker was also inserted to select for the targeted integration. It is important to note that the two copies of *ura4* are in opposite orientations, ensuring that recombination must be driven by the overlap in homology within *kanR* without contributions of the *ura4* sequence. In this assay, use of the homoeologous template during HR allows for the reconstitution of a functional Kan-resistance marker. In WT cells we find that the rate of acquiring Kan-resistance is relatively rare, with a rate of ∼2.1 x 10^-8^ (Fig. 5C and Supplemental Table S1). We found that loss of either *kms1* or *cbp1* causes a significant increase in reconstituting a functional *kanR* cassette, although we observe higher variation in the *cbp1Δ* genetic background (6.24 x 10^-8^ for *kms111*, and 6.03 x 10^-8^ for *cbp111*)(Fig. 5C and Supplemental Table 1). This observation is consistent with tethering to the nuclear envelope (via Cbp1) and/or association with the LINC complex contributing to stringency in selecting the ideal template during repair by homologous recombination.

To provide additional insight into the DNA repair mechanism(s) at play, we further investigated the products of DNA repair leading to acquired Kan-resistance. Using PCR primers on either side of *pfl10* or *pfl4* and both sides of the *kanR*, we determined losses and/or gains of genetic material (Fig. 5D). The *kanR* cassette fragments were retained at the starting loci (“product #1” and “product #4”) in all conditions tested, arguing against a crossover product. In five representative Kan+ WT isolates we detected no change in the structure at *pfl10* whereas there was a gain of the reconstituted *kanR* cassette at *pfl4*, which is indicated by the gain of a PCR product (“product #3”) between the 5’ region of the *kanR* and the flanking *pfl4* (unique) coding sequence (Fig. 5D). The simplest explanation for this result is that a spontaneous DSB at *pfl4* led to synthesis-dependent strand-annealing (SDSA) repair event using *pfl10* as a template, resulting in gene conversion. Kan+ isolates acquired in the absence of Cbp1 similarly led to the gain of the *kanR* cassette at *pfl4*. Of note, in one of five examples we tested we also observed a product of the 3’ region of the *kanR* at *pfl10* (“product #2”), suggesting that new DNA synthesis occurred at both *pfl4* and *pfl10* leading to reconstitution of an intact *kanR* cassette at both loci. Surprisingly, in five representative *kms1Δ* Kan+ isolates we detected no change at *pfl4* and instead a gain of the 3’ region of *kanR* at *pfl10*, suggestive of repair of a DSB at *pfl10* by SDSA using *pfl4* as a template. These observations suggest that although all genotypes reconstitute *kanR* primarily through SDSA, either the spontaneous DSBs driving the repair event and/or their preferred templates are different.

### Kms1 antagonizes mutagenic repair

We previously observed that Kms1 acts as a suppressor of the growth defect observed in HR-deficient *rad5111* cells both on rich medium and in the presence of exogenous DNA damage (Swartz, 2014). However, the mechanism underlying this observation has remained enigmatic. We first sought to test if the improved growth reflects an amelioration of DNA damage load. To this end, we monitored spontaneous DNA repair foci by visualizing Rad52-GFP (Figs. 6A-B). We observed that ∼50% of *rad5111* cells had at least one Rad52-GFP focus at steady state (Figs. 6A-B). While loss of Kms1 had little effect in its own, consistent with our prior report (Swartz, 2014), its deletion in the context of the *rad5111* background led to a marked decrease in spontaneous Rad52-GFP foci to ∼25-30% of cells. Interestingly, most WT or *kms111* cells that contained a Rad52 focus were likely in S phase of the cell cycle (indicated by cell length and morphology), while in *rad5111* or *kms111 rad5111* cells Rad52-GFP foci were prevalent in the G2 phase when cells are quite long, indicating that the damage is persistent (Fig. 6B, transmitted light images). Taken together, these results support the idea that loss of Kms1 allows HR-deficient cells to survive by upregulating another “salvage” DNA repair pathway, as we previously speculated (Swartz, 2014). A pathway able to promote cell survival in the absence of Rad51 in mitotically growing *S. pombe* would be expected to be capable of acting downstream of DSB resection, as essentially all DSBs are committed to HR in this model (Langerak et al., 2011; Leland et al., 2018).

**Figure 6.**
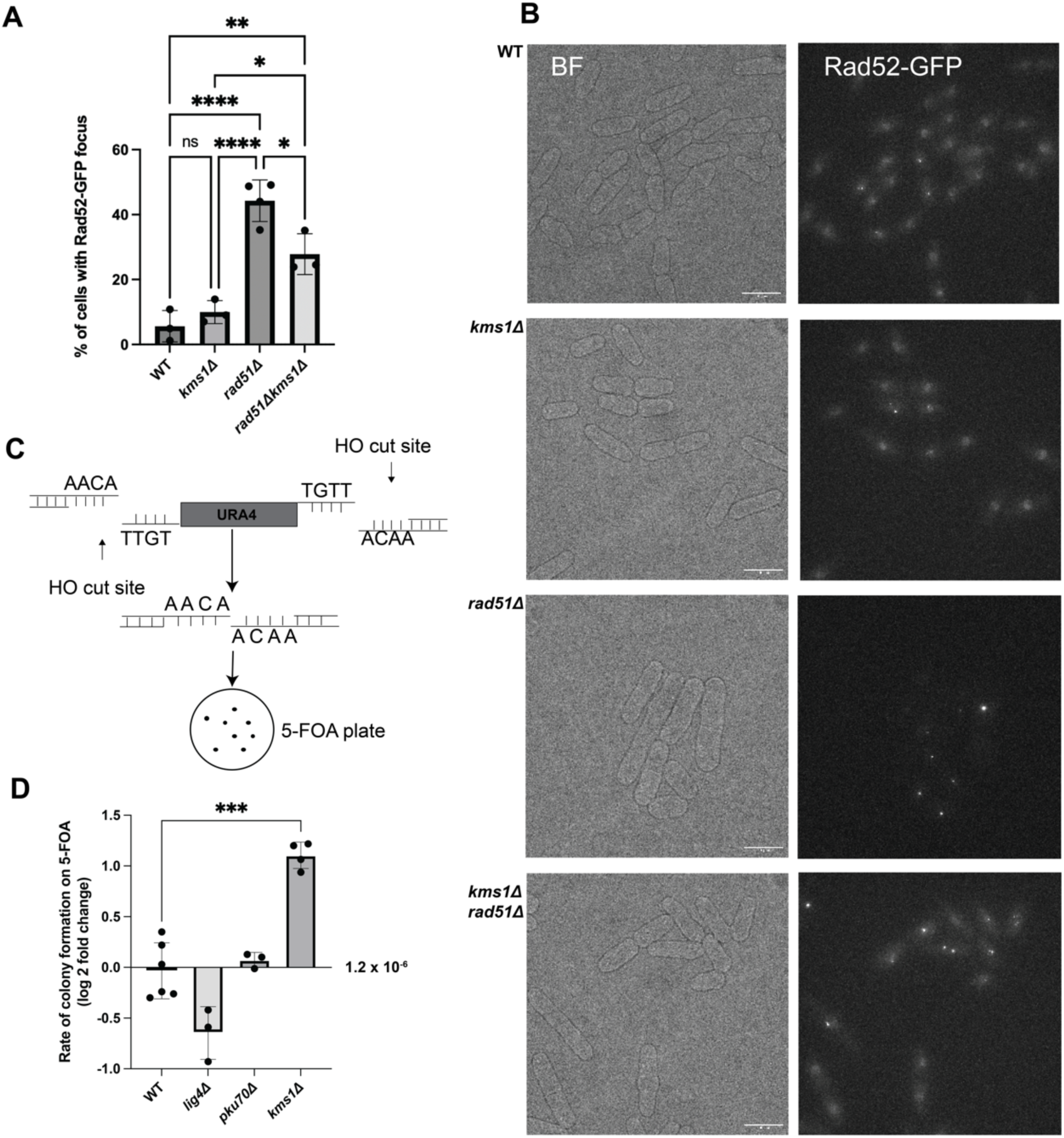
Disrupting *kms1* upregulates mutagenic DSB repair. A) Analysis of Rad52-GFP foci. ∼50% of *rad51Δ* cells have a Rad52-GFP focus. When *kms1* is also lost from a *rad51Δ* background, the number of foci decreases to ∼25-30% (p=0.0158 by one way ANOVA with Tukey correction for multiple comparisons). B) Representative micrographs of Rad52-GFP foci in WT, *kms1Δ*, *rad51Δ*, and *kms1Δrad51Δ* cells. C) The assay to measure the rate of alternative repair pathway usage. D) Loss of the cNHEJ ligase Lig4 causes a trend of a decrease in the rate of loss of *ura4*, suggesting Lig4 is involved in this pathway. Although this finding approached statistical significance on an ANOVA with a Dunnett correction for multiple comparisons (p=0.053), a pairwise t-test revealed a p value of p=0.0151. Loss of *pku70* has no effect, indicating that protection of DSB ends is dispensable in this context. However, loss of *kms1* leads to an increase in the rate of loss of *ura4* (p<0.0001 by one-way ANOVA with Dunnett correction for multiple comparisons), indicating that Kms1 restrains the use of this alternative pathway. Data points are the natural log of the fold change in rate.). * indicates p < .05; ** indicates p < .01; *** indicates p < .001; **** indicates p<.0001

To further investigate this hypothesis, we sought to develop an assay to measure the rate of error-prone DNA repair by creating a pair of inducible DSBs that create incompatible ends, an approach previously used to reveal DNA repair by microhomology-mediated end-joining (MMEJ, also known as alternative end-joining) in budding yeast (Ma et al., 2003b)(Fig. 6C). Specifically, we inserted a *ura4* cassette flanked on both sides by an HO endonuclease cut site into the *mmf1* locus of *S. pombe*, a genomic site that we have previously characterized (Leland et al., 2018) and which resides in the nuclear interior (Zhao et al., 2016). The two HO cut sites are in opposite orientations, giving rise to incompatible DSB ends that are expected to disfavor repair by cNHEJ. As the assay is built in a haploid strain lacking a template for HR, this arrangement essentially forces the cell to use an alternative mechanism to repair the induced DSB (or to die). Therefore, measuring the rate by which viable progeny have lost functional *ura4* allows us to calculate the frequency of alternative repair pathway usage. We first explored the sensitivity of the assay to disrupting cNHEJ. Relative to the rate of colony formation on 5-FOA for WT cells of 1×10^-6^, a *lig4Δ* genetic background led to a ∼50% decrease (Fig. 6D). This raises the possibility that this ligase, typically associated with cNHEJ, contributes to a subset of repaired products that have lost *ura4*. However, we note that loss of the cNHEJ factor Pku70 has no effect, consistent with the expectation that the protection of DSB ends, irrelevant in this engineered system, does not play a role. Consistent with the hypothesis that Kms1 antagonizes an alternative repair mechanism, we observed a three-fold increase in the rate of 5-FOA-resistant colonies in the *kms1Δ* background (Fig. 6D). Taken together, these observations support the idea that the LINC complex restrains cells from using an alternative repair pathway that drives the loss of *ura4*.

### Kms1 restrains the use of an alternative, microhomology-driven repair pathway

One pathway that is important for the survival of HR-deficient cells across model systems is MMEJ (Decottignies, 2007; Lee & Lee, 2007; Sfeir et al., 2024; Truong et al., 2013). Although the fission yeast genome, like budding yeast, does not encode a polymerase with the same structure as the mammalian Pol-θ/PolQ, the use of microhomology upstream of an end joining reaction was first documented in budding yeast (Ma et al., 2003a). We therefore took a sequencing approach to shed light on the type of repair products produced in this alternative repair assay in WT cells and cells lacking *kms1* or *lig4*. To that end, we interrogated individual 5-FOA-resistant isolates from all three genotypes and used colony PCR to amplify across the predicted junction of the two HO cut sites (Fig. 7A). To control for technical failure of the colony PCR for possible large products, primers producing a second, 3.5kb control product were included. When using primers annealing ∼250 bps outside both sides of the HO cut site-flanked *ura4* insertion we found that WT cells mainly produce a small PCR product of the expected size resulting from loss of the entire sequence between the two HO cut sites (∼500bp), although in two of the ten cases (Fig. 7B, red asterisks) we observed a slightly larger product (further discussed below). The 3.5 kb control product is not produced as the smaller products dominate in the PCR reaction. Interestingly, although the *kms1Δ* background led to six of ten representative products of a similar size to those observed in WT cells, we also observed larger products in some *kms1Δ* isolates, suggesting that *kms1Δ* cells can produce a different type of repair product than WT cells. Similarly, *lig4Δ* cells failed entirely to produce the products that dominate in WT cells, in line with a decrease in overall rate (Fig. 6D), instead revealing a surprising preference for exclusively large repair products (Fig. 7B). These results raise the possibility that cells lacking *kms1* or *lig4* can access a similar pathway that is disfavored in WT cells.

**Figure 7.**
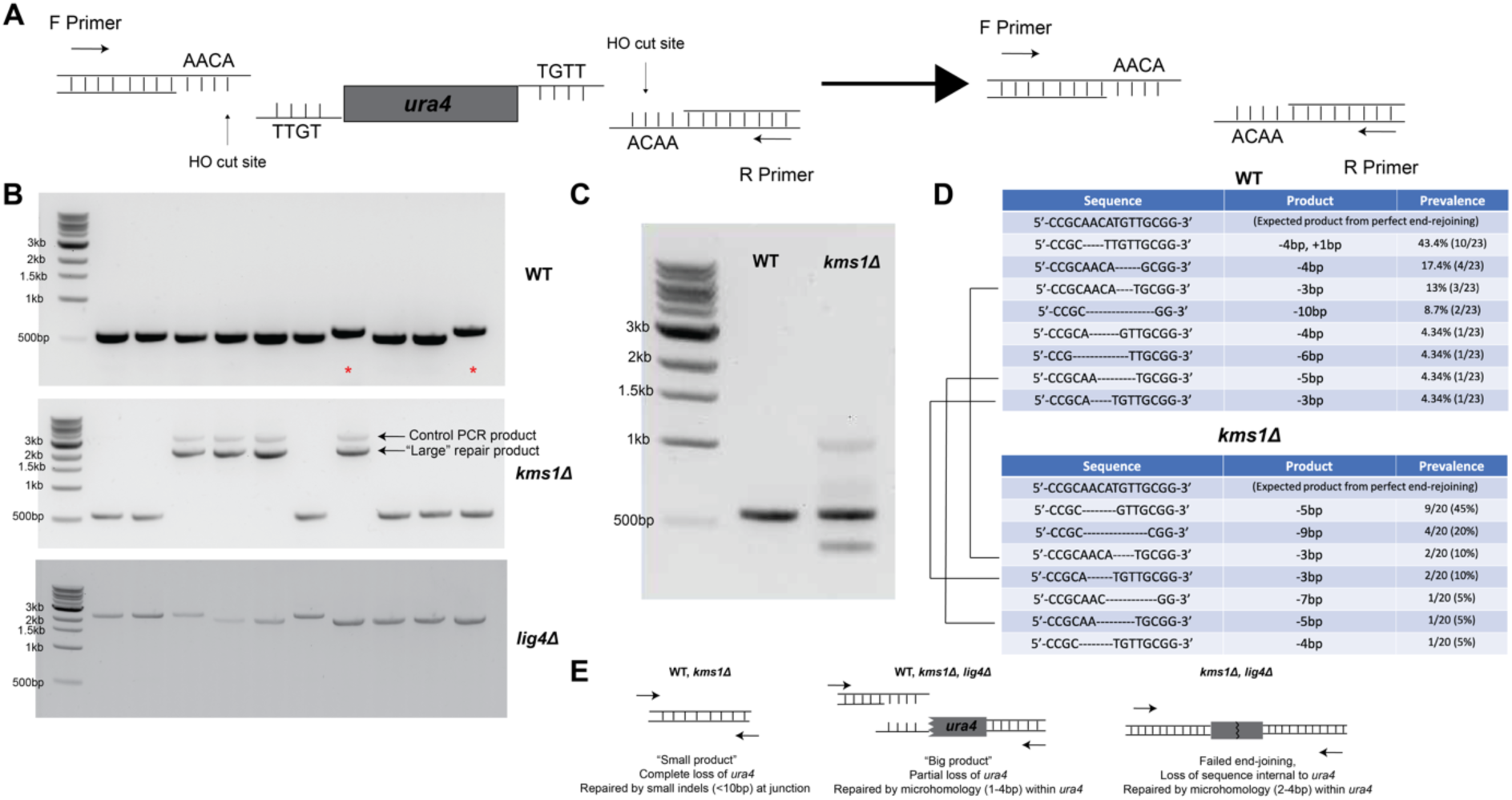
Kms1 restrains the use of a microhomology-driven DNA repair pathway. A) Analysis approach for 5-FOA-resistant colonies. B) PCR amplifying across the junction of the HO sites produces products of the expected size (∼500bp) resulting from loss of the entire sequence between the two cut sites with slightly larger products also observed (red asterisks) in WT cells, while *kms1Δ* cells produce both WT-like small products and larger products, which are similar to products observed in the *lig4Δ* background. To control for technical failure of the colony PCR, a 3.5kb “control product” was also generated. C) PCR analysis of a pool of genomic DNA confirmed that *kms1Δ* cells can produce a mixture of products. D) Sequencing reveals that the most common product in WT cells is produced by a 4bp overhang left by the HO site 5’ to *ura4*, and the untemplated insertion of a single T base. In *kms1Δ* cells producing a similarly sized product to WT cells there are small losses of sequence at the junction but no untemplated insertions. Three products were in common (black bars). E) WT and *kms1Δ* cells produce isolates that result from cleavage of both HO sites and lead to small deletions/insertions at the DSB junction. Larger products in *kms1Δ* cells are produced when the 3’ site and a portion of the *ura4* sequence is retained through use of a microhomology within the *ura4* gene body. In the *lig4Δ* background, neither end of the DSB is able to simply rejoin, and instead repair using microhomology within the *ura4* gene leads to loss of a portion of the sequence. These products are also observed in *kms1Δ* cells, further supporting the idea that *kms1Δ* cells can produce a variety of products.

To further confirm these observations on a greater number of isolates, we prepared a pool of genomic DNA from entire 5-FOA plates of colonies (ranging in number from 30 to 100). With the caveat that smaller products will dominate in the PCR amplification of this pool, we generally confirmed our finding that WT isolates produce expected products in the range of ∼500 bps, while *kms1Δ* isolates produce a mixture of WT-like products and products of both larger and smaller sizes (Fig. 7C), suggesting greater flexibility in the types of repair events that lead to viable progeny that no longer express functional *ura4*. To provide greater insight into how loss of *kms1* affects the repair products (likely contributing to the increase in the rate of forming 5-FOA-resistance colonies, Fig. 6D), we sequenced across the DSB junction in individual isolates that grew on 5-FOA and cloned and sequenced PCR products produced from the pooled genomic DNA. Sequencing of >20 individual isolates revealed that the dominant product in the WT strain (∼50%) lost the 4bp overhang left by the HO site 5’ to *ura4*, followed by an untemplated insertion of a single T base (Fig. 7D), a hallmark typically ascribed to cNHEJ (Decottignies, 2007; Manolis et al., 2001). Notably, a similar chromosomal DSB repair assay in fission yeast revealed a single A base insertion in the dominant repair product in WT cells, which was found to be dependent on Lig4 (Li et al., 2012). The remainder of WT colonies exhibited diverse, small losses of sequence ranging from 3bp to 10bp (Fig. 7D). Given that most products in WT cells do not seem to involve microhomology (Supplemental Fig. 6A), the rate of producing 5-FOA-resistant colonies is lower in *lig4Δ* cells (Fig. 6D), and these repair outcomes were not observed in cells lacking *lig4* (Fig. 7B), the predominant mechanism at play in WT cells appears to be consistent with a cNHEJ-like mechanism (Fig. 7D).

Focusing first on *kms1Δ* isolates that produce a similarly sized product to WT, we observed small losses of sequence at the junction of the two HO sites, but notably no untemplated insertions that resemble the dominant, cNHEJ-like product in WT cells (Fig. 7D). Indeed, we only observed rare products in common between WT and *kms1Δ* isolates (Fig. 7D, black lines), further suggesting that cells lacking Kms1 might engage a distinct mechanism that is less like cNHEJ. To further characterize the larger (>2kb) repair product in *kms1Δ* cells, we cloned and sequenced the dominant species. Interestingly, these products appeared to result from the cleavage of only one of the two HO recognition sites. While the HO site 3’ to *ura4* remains intact, the site 5’ to *ura4* is destroyed. The junction appeared to result from rejoining to microhomology within the gene body of *ura4*, leading to loss of hundreds of base pairs of sequence (Supplemental Fig. S7B). These cells therefore retain a portion of the *ura4* sequence but become resistant to 5-FOA due to the non-functional protein product. The slightly larger WT products (Fig. 7B, asterisks) are also the result of cleavage of only one HO cut site, but notably the only “single cutter” product in WT results in near complete loss of the *ura4* sequence, as opposed to *kms1Δ* cells where a greater span of the *ura4* gene is maintained. This result suggests that cells lacking *kms1* are more able to use microhomologies within *ura4* to drive repair when a single HO cut site is cut, consistent with the elevated overall rate of generating *ura4*-deficient colonies (Fig. 6D). Interestingly, sequencing across the DSB junction in *lig4Δ* cells revealed that both HO cut sites were intact in all samples, suggesting that either “spontaneous” DSBs within the *ura4* were repaired in an error-prone fashion (while those cut at the HO sites could not be repaired and were therefore lost due to lethality) or that DSBs at the HO cut sites were repaired efficiently by another ligase. Interestingly, however, products from *lig4Δ* cells also showed a loss of sequence at *ura4* similar to the *kms1Δ* “single cutter” isolates. These DSBs were repaired by a 4 bp microhomology within the *ura4* gene body (Supplemental Fig. S7C), causing the loss of hundreds of base pairs of sequence internal to *ura4* and a non-functional protein product. We disfavor the interpretation that these products are truly “spontaneous”, however, because we failed to recover any colonies on 5-FOA in the absence of DSB induction. Similar products were also detected in *kms1Δ* cells in which repair proceeded by a 2 bp microhomology (Supplemental Fig. S7B), indicating that this type of repair is also possible when *lig4* is present, although end-joining using the HO site still dominates in this context. Importantly, we never observed the retention of both HO cut sites in WT cells. Taken together, these observations suggest that Kms1 (and therefore a functional LINC complex) normally antagonizes repair by Lig4-independent mutagenic end joining in fission yeast (Fig. 7E).

## Discussion

The fidelity of homology-directed repair depends on the cell’s ability to select the correct homologous template, which can be particularly challenging for DSBs that occur in repetitive regions of the genome. In this work, we leveraged the *S. pombe* flocculin-like gene family as a model to study mechanisms that contribute to genome integrity in repetitive coding genes, as their modular sequences make them prone to instability and intragenic recombination. We identified two factors that influence the fidelity of homology-directed repair outcomes in these gene family: 1) the CENP-B like protein Cbp1, which constrains the *pfl* genes to the nuclear periphery and enforces their stability, and 2) the LINC complex component Kms1, which connects DSBs at the nuclear periphery to dynamics driven by the cytoplasmic microtubule cytoskeleton and disfavors use of homoeologous templates or microhomology-mediated repair. Our observations provide evidence that genome organization contributes to the stability of protein repeat-encoding genes, specifically through constitutive association with the nuclear periphery, while Kms1 enforces stringency in homology-directed repair and antagonizes use of microhomologies, potentially for any DSB that associates with the LINC complex.

### Stability of the *pfl* genes is tied to *Tf2* transposons, Cbp1, and the nuclear periphery

Our findings extend prior studies of genome stability within heterochromatic repetitive DNA elements such as centromeres and telomeres by supporting the idea that tethering of repetitive protein coding genes to the nuclear periphery likewise protects their stability. We demonstrated that most *pfl* genes constitutively reside at the nuclear periphery through a mechanism involving the *Tf2* transposons and/or their LTRs and the LTR-binding factor, Cbp1 (Figs. 1, 3, and 4). We provide evidence that loss of Cbp1 leads to untethering of *pfl5* from the nuclear periphery that we tie to an increase intragenic recombination (Fig. 4); loss of Cbp1 likewise enhances the instability of *pfl3* (Fig. 4). While we observe a more muted effect of abrogating Cbp1 on the position and stability of *pfl10*, we observe that deletion of the nearby *Tf2-3* transposon also enhances repeat instability (Fig. 4). Taken together, these observations suggest that the stability of members of the *pfl* gene family could be coupled, by virtue of their proximity to transposable elements or their remnants (LTRs), to the biology of transposons. As *Tf2* elements are known to cluster in the nucleus in a mechanism that suppresses their mobilization (Cam et al., 2008; Lorenz et al., 2012; Murton et al., 2016), and there is evidence that propagation of the *Tf2* transposons in *S. pombe* rely on HR (Hoff et al., 1998), one attractive hypothesis is that *S. pombe* leverages adaptive mechanisms to restrain transposon activity to stabilize host repetitive coding genes. Indeed, a recent study provides evidence that transposons in *S. pombe* contribute to the fitness of the “host” (Cranz-Mileva et al., 2024). In addition to reducing intragenic recombination of the *pfl* genes, we also observe that Cbp1 restrains intergenic recombination between the related *pfl4* and *pfl10* (Figs. 5C-D). Cbp1 might also further restrain the likelihood that two *pfl* genes can encounter one another after a spontaneous DSB arises by virtue of tethering them to the nuclear periphery, as the probability of a physical encounter between the DSB and template is thought to play a strong contribution to template choice during HR (Wang et al., 2017).

### Kms1 enforces the fidelity of homology-directed repair

Most *pfl* genes reside at the nuclear periphery constitutively (Fig. 1), but we also previously demonstrated that slow-to-repair DSBs arising in the nuclear interior are targeted to Sad1-Kms1 LINC complexes in fission (Swartz, 2014). The stability of *pfl5* is dominated by the effect of Cbp1, perhaps because it is constitutively compartmentalized at the nuclear periphery. However, we observe that loss of Kms1 enhances the likelihood of an intergenic repair event between *pfl4* and *pfl10* (Fig. 5C) and also alters the recombination products (Fig. 5D). It is not yet clear why cells lacking Kms1 prefer to reconstitute *kanR* at *pfl10* rather than *pfl4*, which dominates in WT cells. One possibility is that this result reflects a more prominent role for Kms1 in antagonizing this homoeologous strand invasion event occurring between a spontaneous DSB at *pfl10* and the donor sequence at *pfl4*. One attractive model is that LINC complex dynamics promote the dissolution of stand invasion structures without sufficient homology in a kinetic proofreading-type mechanism, thereby enforcing stringency in template choice during any type of homology-directed repair. However, it is also plausible that the loss of Kms1 influences the relative likelihood that a spontaneous DSB arises at these two loci.

### The LINC complex restrains mutagenic MMEJ

Consistent with a physical model in which Kms1 promotes dissolution of repair intermediates lacking sufficient homology, at an irreparable DSB Kms1 appears to antagonize MMEJ, as we observe up-regulation of repair events leveraging microhomologies in *kms1Δ* cells (Fig. 6D; Fig. 7B-E, Supplemental Fig. S7). This observation provides an explanation for our previous finding that loss of Kms1 can suppress the growth defect of *rad51Δ* cells (Swartz, 2014). Accordingly, loss of Kms1 decreases the frequency of spontaneous Rad52 foci in *rad51Δ* cells (Figs. 6A-B). These results support the idea that a Rad51-independent repair mechanism is up-regulated upon loss of fully-assembled LINC complexes. Using a system employing two inverted site-specific DSBs to enforce incompatible ends (Fig. 6C), we observe that WT cells repaired primarily with an untemplated insertion of a single T base at the junction of the two DSB ends (Fig. 7D), similar to a prior study that reported a frequent single untemplated A base insertion product that was lost in the absence of *lig4* (Li et al., 2012), reinforcing the idea that this reflects a cNHEJ product.

Our inability to detect the dominant, WT cell product in *lig4Δ* cells further supports this interpretation. Instead, we observe that loss of *lig4* leads to an alternative repair outcome that drives an internal deletion in *ura4* with junctions containing a 4 bp microhomology (Supplemental Fig. S7). Along with greater proficiency in repairing these incompatible DSB ends, *kms1Δ* cells also disfavor the cNHEJ-like single base pair insertion (Fig. 7D). Instead, we observe a greater variety of repair products of varying sizes that result from microhomology within the *ura4* gene body (Figs. 7B-7C and Supplemental Figs. S7B-C). Taken together, these results support the idea that the LINC complex restrains the use of repair by the mutagenic, *lig4*-independent, MMEJ pathway, which cells will use when recombination with a homologous template is not possible.

Overall, our findings propose two mechanisms that enforce the maintenance of genome integrity in fission yeast, particularly in repetitive regions of the genome. First, we find that tethering repetitive, protein-coding genes to the nuclear envelope via the transposon-binding protein, Cbp1 restrains both intragenic recombination and a non-allelic recombination event between homoeologous genes. Kms1, the cytoplasmic aspect of the LINC complex relevant to DSB repair in fission yeast, similarly represses illegitimate recombination events within and between repeat-encoding genes as well as mutagenic MMEJ when recombination is inhibited. These mechanisms contribute to the broad evidence that nuclear compartmentalization helps to ensure faithful DNA repair.

## Materials and Methods

### Microscopy

For lacO localization, images were acquired on a DeltaVision wide-field microscope (Applied Precision/GE) using a 1.2 NA 100x objective (Olympus), solid-state illumination, and an Evolve 512 EMCCD camera (Photometrics). Cells were mounted on 1.2% agar pads and sealed with VALAP (1:1:1 vaseline:lanolin:paraffin). Imaging parameters were as follows. Transmitted light: 50% transmittance, 0.025s exposure; mCherry: 50% power, 0.1s exposure; GFP: 50% power, 0.01s exposure. 40 Z-sections were acquired at 0.26mm spacing. For spontaneous Rad52 foci, images were acquired on the same microscope and objective but with a CoolSnap CCD camera. Cells were mounted directly on a glass slide. Imaging parameters were as follows: Transmitted light: 32% transmittance, 0.015s exposure; mCherry: 32% power, 0.1s exposure; GFP: 32% power, 0.1s exposure. 16 Z-sections were acquired at 0.4mm spacing.

### 3D particle localization and image analysis for 3D nuclear reconstruction

We determined the position of GFP-labeled genomic loci from a set of 2D microscope images collected at multiple equally-spaced focal depths (z-stack). To estimate the 2D positions within the plane of the microscope field of view, we generate a maximum intensity projection over all 2D images (z-slices). We then estimate the 2D positions of each locus to pixel level accuracy by thresholding and keeping only the local maximum intensity within a given radius of 1.5 pixels, which is sufficient to encompass each locus. We then select a region around the estimated locus position from the full z-stack and perform a least-mean-squares fitting of a 3D Gaussian within the selected region to determine the best-fit position of locus. We used a previously developed image analysis pipeline outlined in our previous work (Zhao et al., 2016). The GitHub folder with the code can be found here: https://github.com/mochrielab/3DMembraneReconstruction. In this pipeline, we first generate an initial spherical mesh for each nucleus with a radius calculated based on the nuclear area of the 2D projection of the z-stack. We then modify the vertices of the mesh to minimize the total energy, which consists of the bending energy of the surface and an energy cost related to how well the shape describes images in the entire z-stack. The reconstruction script also produces nuclear volume measurements (Supplemental Fig. S5A).

### Probability density calculation

We calculated the probability densities for Figs. 1 D and F, Figs. 4 C-E, and Supplemental Fig. S4 as follows. First, we scaled the coordinates for each cell by the corresponding nuclear radius. Then, we partitioned each scaled nucleus into concentric shells of width, 𝑤, equal to 0.1 of the scaled nuclear radius. Next, we determined the probability, 𝑝, that each locus of interest is observed in the shell at scaled radial coordinate, 𝑟. Then, the probability density, 𝑓, is 𝑝 divided by the scaled volume of the corresponding shell: 𝑓 = 𝑝⁄(4𝜋𝑟^2^𝑤). The scaled distance from the nuclear envelope, 𝑠, is the complement of the scaled radial coordinate, i.e. s = 1 − r. We then plot 𝑓 = 𝑝⁄(4𝜋(1 − 𝑠)^2^𝑤) versus 𝑠. Since 𝑝, 𝑟, 𝑠, and 𝑤 are all dimensionless, so is the probability density in this case. The error bars represent the Poisson counting uncertainty propagated to the probability density. For a bin with c points (out of a total of N), the uncertainty is 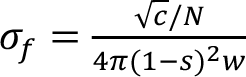.

### Yeast culture and strain construction

*S. pombe* cells were grown, maintained, and crossed using standard procedures and media (Moreno et al., 1991). Gene replacements were made by gene replacement with various MX6-based drug resistance genes (Bähler et al., 1998; Hentges et al., 2005). For lacO strains: plasmids previously designed in our group, pSR10_ura4_5.6kb that contain a series of lacO repeats 5.6kb in length (Leland & King, 2014) were targeted to various *pfl* genes using the primers listed in Supplemental Table S4. For intragenic and intergenic/homoeologous recombination strains: megaprimers were amplified from pFA6a-Kan that contained opposite halves of the Kanamycin-resistance gene (with ∼220bp of overlap between them), with homology arms to target *ura4* in the plasmid pAW1. These megaprimers were used to amplify *ura4* from plasmid pAW1 using iProof polymerase, creating a cassette that contained both halves of the Kanamycin-resistance gene and *ura4*. This cassette was targeted to various *pfl* genes (*pfl3*, *pfl4*, *pfl5*, *pfl*10, *mam3*, *map4*) of *S. pombe*. For the “incompatible DSB ends” DNA repair assay strains: pFA6a_HOcs_Ura_HOcs was integrated into the *mmf1* locus of *S. pombe* using the primers Mmf1L1 and Mmf1L6 listed in Supplemental Table S4.

### Plasmid construction

To produce pFA6a_HOcs_Ura_HOcs we modified a plasmid from a previous study (Leland et al., 2018) that contains a modified MX6-based HygR cassette containing the HO cut site (pFA6a-HOcs-Hyg). We first swapped the marker to the *S. pombe ura4* gene by cutting at AscI and PmeI to release the HygR fragment followed by inserting the *S. pombe ura4* gene amplified from a pFA6a-*ura4* plasmid with primers containing AscI and PmeI cut sites. Next, the second HO cut site was inserted into pFA6a_HOcs_Ura on the opposite side of the *ura4* insert in the opposite orientation of the existing HO cut site by cutting at AscI and annealing the following complementary, phosphorylated oligonucleotides containing AscI cut sites on both ends: HOcs_ins_AscI_F and HO_ins_AscI_R to produce pFA6a_HOcs_Ura_HOcs. The HOcs_Ura_HOcs cassette was amplified using iProof polymerase-based PCR with the primers Mmf1L1+Mmf1L6. The cassette was transformed into the *mmf1* locus of *S. pombe* using standard procedures.

### Assay for measuring instability at repetitive regions of *pfl* family genes

Single colonies were picked from strains maintained on EMM-Ura medium and grown to saturation in 5mL YE5S medium at 30 °C for 24-48 hours. Cultures were plated to 5-FOA (diluted 1:100) and YE5S (diluted 1:10,000,000). After 3 days at 32 °C, the colonies on both plates were counted to calculate the rate of loss of functional *ura4*. Fold change in rate was calculated by dividing each individual rate by the mean rate in WT. All plots show the natural log of the fold change in rate. For over-expression using the *nmt41* promoter: single colonies from EMM-Ura+Thi (+Thi to repress the *nmt41* promoter) were grown to saturation in 5mL EMM-Ura medium with or without thiamine at 30 °C for 48 hours. Cultures were plated to 5-FOA and YE5S as described above.

### RT-qPCR to measure expression of *pfl* family genes

Cells maintained on EMM-Ura were grown in 5mL YE5S medium at 30 °C for 20 hours. RNA was extracted using the MasterPure Yeast RNA Purification Kit (Lucigen). Reverse transcription was carried out using the Verso cDNA Synthesis Kit (ThermoScientific), along with a blend of random hexamers and anchored oligo-dT 3:1 and qPCR reactions were carried out with 2X iTaq Universal SYBR Green (BioRad) and forward/reverse primers for the target *pfl* gene or for *act1* (Supplemental Table S4). Cq values were analyzed using the ΔΔCq method, with *act1* for normalization with three technical replicates per condition.

### Chromatin immunoprecipitation plots

For H3K4me3, reads from SRR948227.fastq.gz and SRR948235.fastq.gz (DeGennaro et al., 2013) were aligned using bowtie to allow all multiple alignments (bowtie --best) and the linear scale fold enrichment file was generated using MACS2 (callpeak and bdgcmp). For H3K9me3, the file RES15-17-P4-ChIP-br172-H3K9triMeChIP_Run1.realn.bam was indexed using samtools (Shan et al., 2016). For Cbp1, the GSM2150395_peaks_cnagdata_geo.bed.gz is displayed (Daulny et al., 2016).

### Assay for measuring homoeologous recombination

Cells maintained on EMM-Ura were grown in 5mL YE5S medium at 30 °C over the day, then diluted 1:10 (to a total culture volume of 50mL) and allowed to grow at 30 °C overnight. They were then plated to YE5S+Kan and YE5S (diluted 1:6,000,000). After 3 days at 32 °C, the colonies on both plates were counted to calculate the rate of acquiring Kan resistance. Fold change in rate was calculated by dividing each individual rate by the mean rate in WT. Plots show the natural log of the fold change in rate. To amplify the products of homoeologous recombination, gDNA was extracted from Kan+ colonies and subjected to PCR using primers to amplify across both loci and the kanamycin-resistance gene. For the starting products, the following primer pairs were used: PFL10For + KanUpRev (“product #1”) and PFL4Rev + TEFTermFor (“product #4”). For the recombinant products the following primer pairs were used: PFL10Rev + TEFTermFor (“product #2”) and PFL4For+ KanUpRev (“product #3).

### Assay for measuring salvage DNA repair pathway

Cells that were maintained on EMM-Ura were transformed with pREP81HO (Prudden et al., 2003) using the lithium acetate method and plated onto EMM-Ura-Leu+Thi (-Leu to select for uptake of the pREP81HO plasmid, and +Thi to repress the expression of the HO endonuclease under the control of the nmt41 promoter). A subset of colonies from that plate (∼10-30) were grown together in 50mL EMM-Ura-Leu+Thi overnight at 30 °C. The following day, the entire culture was spun down and washed twice with water, diluted in EMM-Leu to an of OD 0.05 and grown at 30 °C over the day. To prevent culture saturation and to allow the cells to continue cycling, cultures were diluted to OD 0.05 at 8 hours, 24 hours, and 32 hours post-thiamine wash. 48 hours post-thiamine wash, the cultures were plated onto 5-FOA and YE5S (diluted 1:2,000,000). After 3 days at 32 °C, the colonies on both plates were counted to calculate the rate of loss of *ura4*. Fold change in rate was calculated by dividing each individual rate by the mean rate in WT. Plots show the natural log of the fold change in rate.

### Sequencing

The junction across the HO cut sites were amplified with flanking primers approximately 250bp outside the insertion (Mmf1L1+Mmf1L6) via colony PCR with 2X Long Amp Taq. An additional primer set (Aly3F and Aly3R in Supplemental Table S4) of primers was used as a control for amplification of large products. For colonies that produced a positive result, genomic DNA was extracted using the phenol/chloroform method and a Taq-polymerase based PCR was performed using the same primer pair for bulk sequencing. After clean up using the Qiagen PCR Purification Kit the products were sequenced priming from the L1Seq oligo (Supplemental Table S4). For “large products”, the purified PCR product was TA cloned into a pCR 2.1 vector using the Invitrogen TOPO TA Cloning Kit, transformed, and plasmid DNA was purified with the Qiagen Miniprep Kit. Plasmids were digested with EcoRI to determine the size of the insert and sequenced.

## Acknowledgements

We would like to thank all members of the LusKing Lab for discussion and feedback, particularly Dr. Connor McBrine. We thank Elisa Rodriguez for technical expertise and support. We also thank the Yeast Genome Resource Center at Osaka University and the many investigators who have deposited strains at this resource. We thank the Keck DNA Sequencing Facility at Yale for their assistance. This work was supported by DP2OD008429 and R35GM153474 (to MCK), EFMA-1830904 (to MCK and SGJM) and T32GM007499 and F31GM143917 (to AL).

## Author Contributions

Conception and design: AL, DL, and MCK.

Development of methodology: AL, DL, NL, YJT, SGJM and MCK.

Acquisition of data: AL and DL.

Analysis and interpretation of data: AL, DL, YJT, NL, SGJM, CPL, and MCK.

Writing original draft: AL and MCK.

Writing, review, and/or revision of the manuscript: AL, DL, YJT, NL, SGJM, CPL, and MCK.

Study supervision: SGJM, CPL, and MCK.

## Notes

### Competing Interest Statement

The authors have declared no competing interest.

### Summary of Updates

Revisions and additional data can be found to the figures: Fig. 1B, Fig. 3C-D, Fig. S3A, There are also new additional figures: Figures S4, S5,

