## Supplemental Figures S1-S7 and Tables S1-S5 for "The LINC complex component Kms1 and CENP-B protein Cbp1 cooperate to enforce faithful homology-directed DNA repair at the nuclear periphery in *S. pombe*"

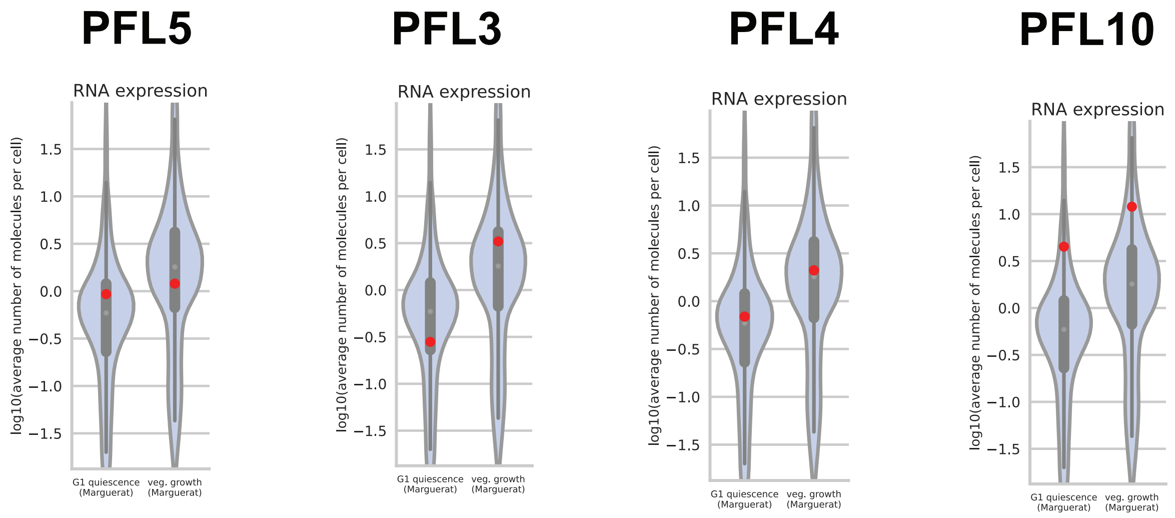

**Supplemental Figure S1. *Pfl* genes are expressed.** Data from a previous study (Marguerat et al., 2012) shows that *pfl*5, *pfl*3, *pfl*4, and *pfl*10 are expressed at a level at or above that of the average gene during the vegetative growth phase.

**SPAPB2C8.01- *pfl*10 repeat amino acid sequence**

| Repeat # | Sequence |
|---|---|
| 1 | NTITTTLYSGSLEFTTTLDSANGTTPATIEVVEPAA |
| 2 | GTVTTTIYSGTTPFNTTLASATDTVPGTVEVVEPEA |
| 3 | GTVTTTVYSGTQEYTTTLATASGTVSGTVEVVDTAA |
| 4 | GTVTTTIYSGTTPFNTTLASATDTVPGTVEVVEPEA |
| 5 | GTVTTTVYSGTQEYTTTLATASGTVSGTVEVVDTAA |
| 6 | GTVTTTIYSGTTPFNTTLASATDTVPGTVEVVEPEA |
| 7 | GTVTTTVYSGTQEYTTTLATASGTVSGTVEVVDTAA |
| 8 | GTVTTTIYSGTTPFNTTLASATDTVPGTVEVVEPEA |
| 9 | GTVTTTVYSGTQEYTTTLATASGTVSGTVEVVDTAA |
| 10 | GTVTTTIYSGTTPFNTTLASATDTVPGTVEVVEPEA |
| 11 | GTVTTTVYSGTQEYTTTLATASGTVSGTVEVVDTAA |
| 12 | GTVTTTIYSGTTPFNTTLASATDTVPGTVEVVEPEA |
| 13 | GTVTTTVYSGTQEYTTTLATASGTVSGTVEVVDTAA |
| 14 | GTVTTTIYSGTTPFNTTLASATDTVPGTVEVVEPEA |
| 15 | GTVTTTVYSGTQEYTTTLATASGTVSGTVEVIEPAA |
| 16 | GTVTTTVYSGTQEYTTTLATASGTVSGTVEVIEPAA |
| 17 | GTVTSTVYSGTQEYTTTLATASGTVSGTVEVIEPAA |
| 18 | GTVTTTVYSGTQEYTTTLATASGTVSGTVEVIEPAA |
| 19 | GTVTTTVYSGSEVYTTTLASASESVPGTVEVVDPEA |
| 20 | GSVTTTIYSGSVEYTTVLADASGSVTGTVEVVEPAA |
| 21 | GTVTTTVYSGSEVYTTTLASASESVPGTVEVVDPEA |
| 22 | GSVTTTIYSGSVEYTTVLADASGSVTGTVEVVEPAA |
| 23 | GTVTGTLTSGSQFFTTTIAQASGSVSGNVEVIEPSG |
| 24 | STVTSTIYSGSESFTTTLAVGSGTIPGTVEVILPAP |

**SPCC188.09C- *pfl*4 repeat mino acid sequence**

| Repeat # | Sequence |
|---|---|
| 1 | STIYTTINSGTVASTYTVTLADGDV--VVKDIEPTA |
| 2 | GTVTTTITSGSHEFTTTLAEASGTVPGTVEVVEPLA |
| 3 | GTVTTTIHSGSVEYNTTLATASGTVPGTVEVVEPAV |
| 4 | GTVTTTIYSGSEEFTTTLASASGSISGTVEVIEPTA |
| 5 | GTVTTTIYSGDQEYTTILAEASGTVPGTVEVIEPAV |
| 6 | GTVTTTTYSGSVEYTTTLVPASGSVSGTVEVVEPAV |
| 7 | GTVTTTLQSGSQAFTTTV-PASGSVSGTVEVVQPTG |
| 8 | GTVTNTVYEGSQTITSTLATASGTVPGTVEVILPGP |
| 9 | STIY----SGTVATTITYDVSSTPASTVVV-I-PTA |

**SPBC1289.15-*pfl*5 repeat amino acid sequence**

| Repeat # | Sequence |
|---|---|
| 1 | GTITETIVSGSVGYTSTFPASGTTSGTVEVVEPTA |
| 2 | GTITETIVSGSVGYTSTFPANGTTSGTVEVVEPTA |
| 3 | GTITETIVSGSVGYTSTFPANGTTSGTVEVVEPTA |
| 4 | GTITETIVSGSVGYTSTFPANGTTSGTVEVVEPTA |
| 5 | GTVTETIVSGSVGYTSTFPASGTTSGTVEVVEPTA |
| 6 | GTITETIVSGSKAFTSTFPANGTTSGTVEVVEPTA |
| 7 | GTITKTIVSGSKTFTSTFPANGTTSGTVEVVEPTA |
| 8 | GTITETIVSGSVGYTSTFPANGTTSGTVEVVEPTA |
| 9 | GTITETIVSGSKTFTSTFPASGTTSGTVEVVEPTA |
| 10 | GTITETIVSGSKAFTSTFPANGTTSGTVEVVEPTA |
| 11 | GTITETIVSGSVGYTSTFPASGTTSGTVEVVEPTA |
| 12 | GTVTETIVSGSVGYTSTFPASGTTSGTVEVVEPTA |
| 13 | GTVTETIVSGSVGYTSTFPASGTTSGTVEVVEPTA |
| 14 | GTITETIVSGSKAFTSTFPANGTTSGTVEVVEPTA |
| 15 | GTITETIVSGSKTFTSTFPANGTTSGTVEVVEPTA |
| 16 | GTITETIVSGSVGYTSTFPASGTTSGTVEVVEPTA |
| 17 | GTVTETVISGSVSFMSTITAHDTSSGAVIVVEPTA |
| 18 | GTVTETIVSGSIPFTSTIPAQGTTSGTVEVVEPTA |
| 19 | GTVTETIISGSVGYTSTFPAQGTTSGTVEVVEPTA |
| 20 | GTVTETIISGSVGYTSTFPAQGTTSGTVEIVAPTA |
| 21 | GTVTETIVSGSIPFTSTIPAQGTTSGTVEVVEPTA |
| 22 | GTVTETIISGSVGYTSTFPAQGTTSGTVEIVAPTA |
| 23 | GTVTETIVSGSIPFTSTIPAQGTTSGTVEVVEPTA |
| 24 | GTVTETIISGSVGYTSTFPAQGTTSGTVEVVEPTA |
| 25 | GTVTETIVSGSIPFTSTIPAQGTTSGTVEIVVPTA |
| 26 | GTVTSTTGSGTSWFTTTVPATGTRSGSVIVVSPTA |

**SPBC947.04-*pfl*3 repeat amino acid sequence**

| Repeat # | Sequence |
|---|---|
| 1 | GTITLTTISGSDLYTSTFPANGTTSGTVEVVIPTA |
| 2 | GTVTETAVSGSELYTSTFPANGTTSGTVEVVIPTA |
| 3 | GTRTVTKISGSKFFTTTTDASGTVSGTVEVVLPTA |
| 4 | GTNTMTVVSGSRFFTSVVSASGTVSGEQVIVYPTA |
| 5 | GMVTETIVSGSEIFNTTYPASGTRTGTVEVVIPTA |
| 6 | GTVTETEISGSELYTSTFPANGTTSGTVEVVIPTA |
| 7 | GTRTVTKISGSKFFTTTTDASGTVSGTVEVVLPTA |
| 8 | GTNTMTVVSGSRFFTSVVSASGTVSGEHIIVEPTA |
| 9 | GVVTETVVSGSVGYTTTYPAHDTVSGTVEVVEPTA |
| 10 | GVVTETVVSGSVGYTTAYPAHDTVSGTVEVVEPTA |
| 11 | GVVTETVVSGSVGYTTTYPAHDTVSGTVEVVEPTA |
| 12 | GVVTETVVSGSVGYTTTYPAHDTVSGTVEVVEPTA |
| 13 | GVVTETVVSGSVGYTTTYPAHDTVSGTVEVVEPTA |
| 14 | GVVTETVVSGSVGYTTTYPAHDTVSGTVEVVEPTA |
| 15 | GVVTETVVSGSVGYTTTYPAHDTVSGTVEVVEPTA |
| 16 | GVVTETVVSGSVGYTTTYPAHDTVSGTVEVVEPTA |
| 17 | GVVTETVVSGSVGYTTTYPVQGTVSGTVMIVEPTA |

**Supplemental Figure S2. *Pfl* genes contain repeats and share sequence homology to each other.** Alignment of the amino acid sequence of *pfl*10, *pfl*4, *pfl*5, and *pfl*3 demonstrate that not only are these genes repetitive, but they also share homology to each other.

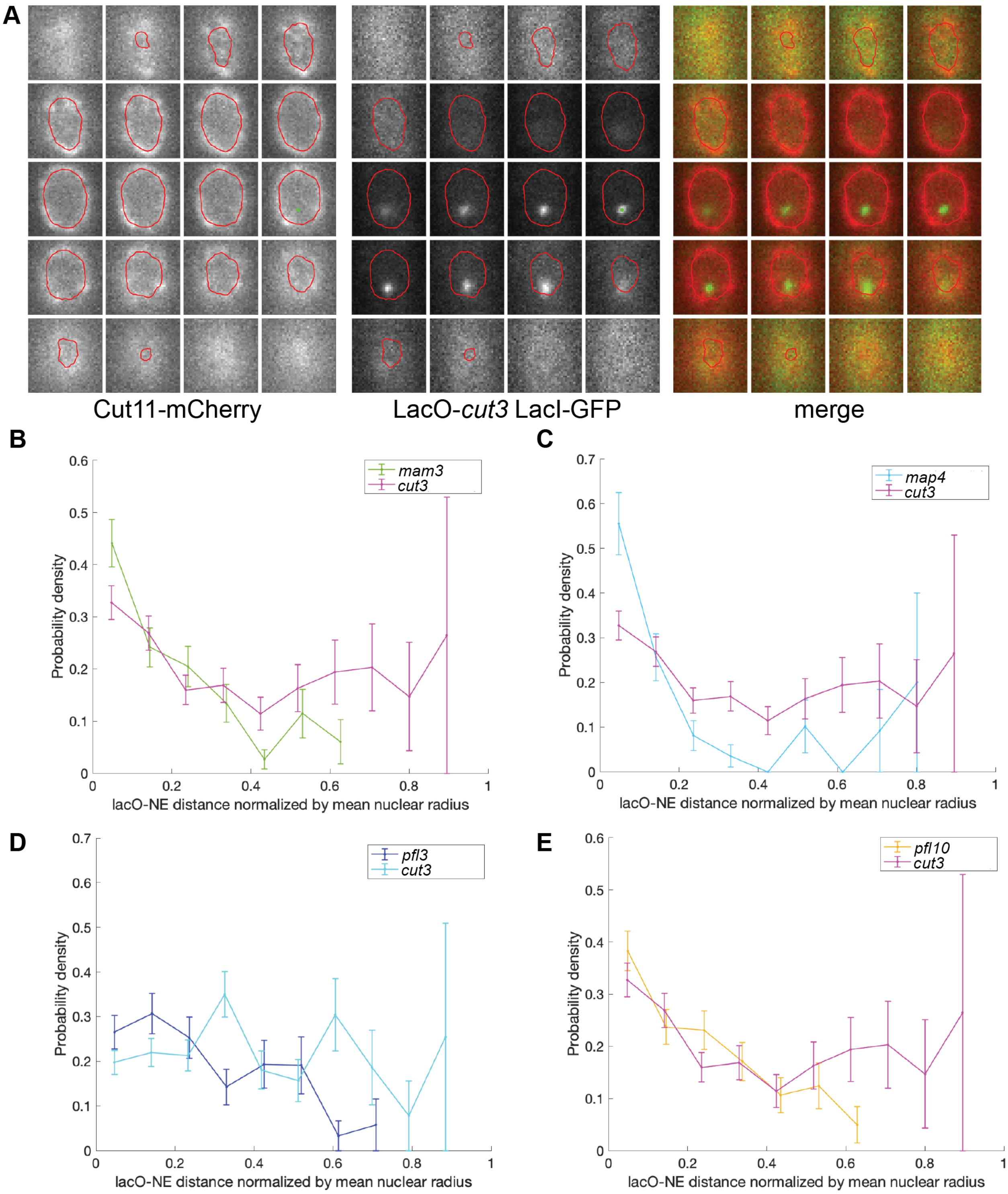

**Supplemental Figure S3. Nuclear reconstruction and *pfl* gene position.** A) Example of the nuclear reconstruction approach showing the Cut11-mCherry signal and fit, lacO position, and the merge. B-E) A subset of *pfl* family genes are enriched at the nuclear periphery, unlike a generic locus, *cut3*. A probability density analysis shows that: B) *pfl10*, C) *mam3*, D) *map4*, and E) *pfl3* prefer to reside close to the nuclear periphery, unlike the generic locus *cut3*.

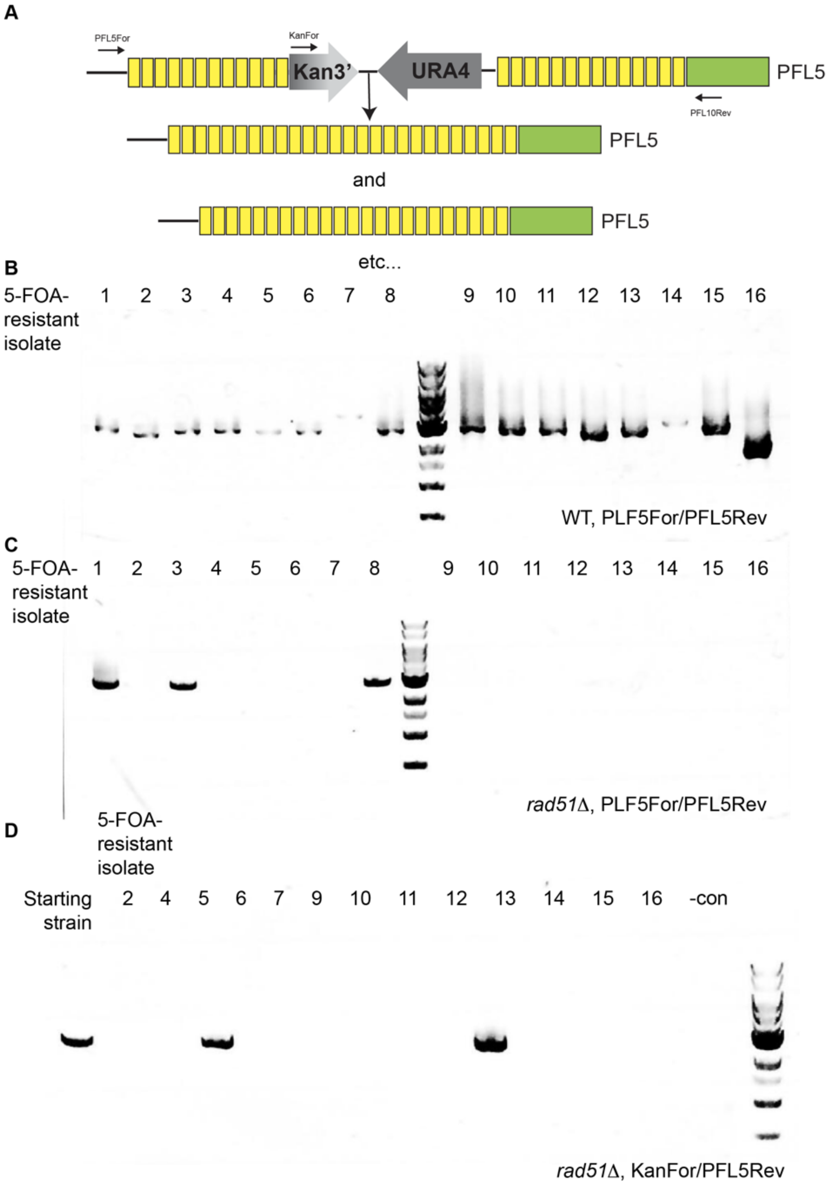

**Supplemental Figure S4. PCR analysis of the 5-FOA resistant isolates.** A) Cartoon of the intragenic assay for loss of *ura4*. B) Products produce by PCR across *plf5* repeats in 5-FOA resistant WT isolates. C) Products produced by PCR across *plf5* repeats in 5-FOA resistant *rad51Δ* isolates. D) Products produced by PCR to detect retention of *ura4* in *rad51Δ* isolates that fail to produce a product in panel (C).

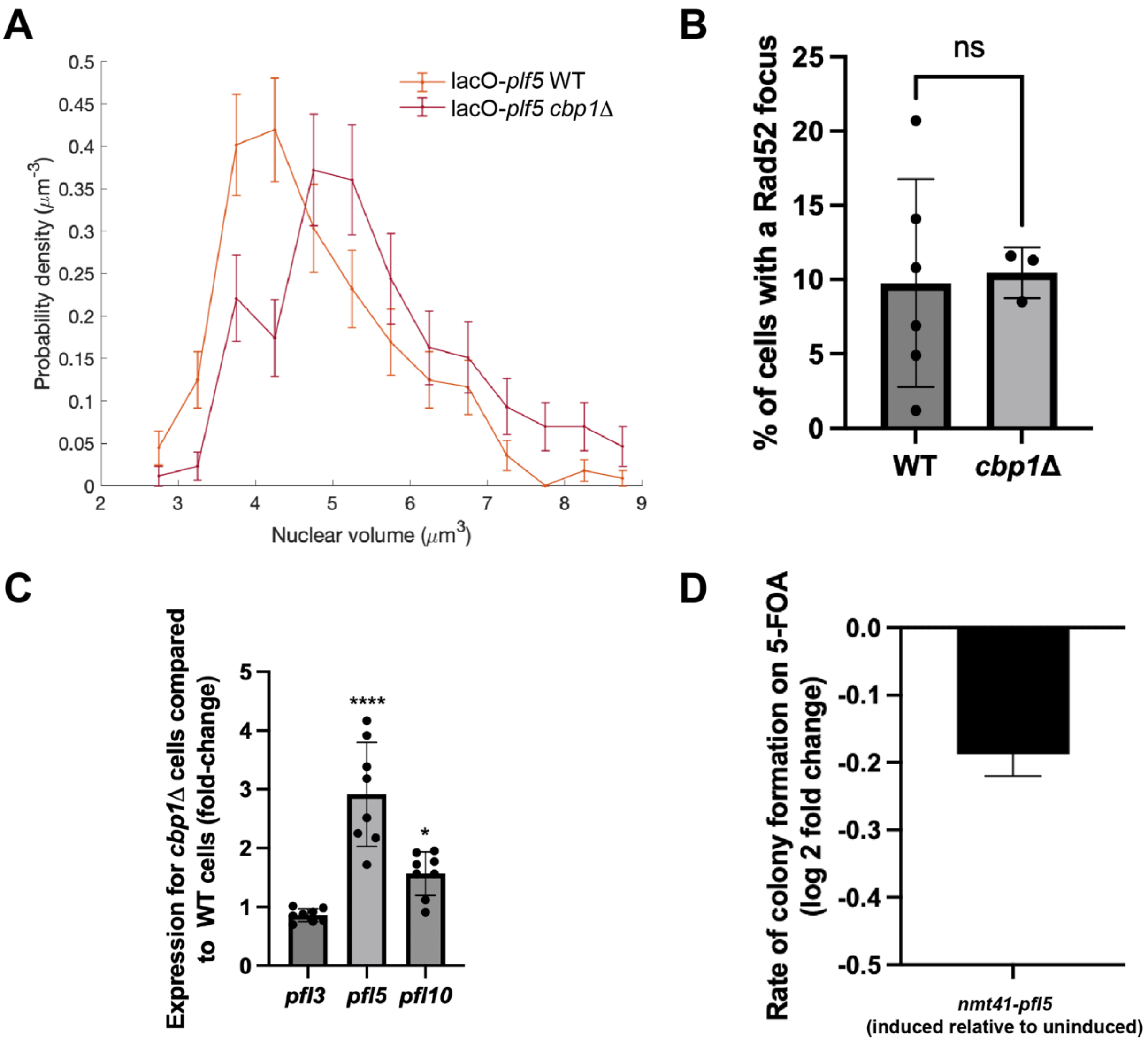

**Supplemental Figure S5. Contribution of Cbp1 to *pfl* gene stability.** A) Cells lacking Cbp1 have subtly (16%) larger nuclear volumes than WT cells. B) Loss of Cbp1 does not affect the prevalence of spontaneous DNA damage as assessed by counting of Rad52-GFP foci. C) Loss of Cbp1 leads to a statistically significant increase in the expression of *pfl5* but not *pfl3* or *pfl10* as assessed by RT-qPCR. Statistical analysis using ANOVA; $p<0.0001$. D) Driving *pfl5* expression by the regulatable *nmt41* promoter does not increase the instability of *pfl5*. Plotted as the log2 fold change of the rate of colonies produced on 5-FOA in the induced (no thiamine) relative to repressed (rich media) condition.

**SPAPB2C8.01- *pfl10* Nucleotide sequence**

| | Sequence |
|---|---|
| 1 | AATACAATAACGACTACTCTTTATTCTGGTAGTCTCGAGTTTACTACTACTCTTGACTCGGCAAATGGGACTACCCCTGCAACCATTGAAGTAGTTGAACCTGCGGCA |
| 2 | GGGACCGTTACCACTACAATTTACTCTGGAACTACACCATTTAACACAACGCTTGCTTCCGCAACTGATACTGTTCCTGGCACTGTTGAGGTAGTTGAGCCTGAAGCT |
| 3 | GGAACTGTCACTACTACTGTATATTCTGGCACTCAAGAGTACACCACAACGCTTGCTACAGCCAGTGGTACTGTTTCAGGCACTGTTGAAGTAGTGGATACAGCAGCT |
| 4 | GGAACTGTTACTACTACAATTTACTCTGGAACTACACCATTTAACACAACGCTTGCTTCCGCAACTGATACTGTTCCTGGCACTGTTGAGGTAGTTGAGCCTGAAGCT |
| 5 | GGAACTGTCACTACTACTGTCTATTCTGGTACTCAAGAGTACACTACCACACTTGCTACAGCCAGTGGTACTGTTTCAGGCACTGTTGAAGTAGTGGATACAGCAGCT |
| 6 | GGAACTGTTACTACTACAATTTACTCTGGAACTACACCATTTAACACAACGCTTGCTTCCGCAACTGATACTGTTCCTGGCACTGTTGAGGTAGTTGAGCCTGAAGCT |
| 7 | GGAACTGTCACTACTACTGTATATTCTGGCACTCAAGAGTACACTACCACACTTGCTACAGCCAGTGGTACTGTTTCAGGCACTGTTGAAGTAGTGGATACAGCAGCT |
| 8 | GGAACTGTTACTACTACAATTTACTCTGGAACTACACCATTTAACACAACGCTTGCTTCCGCAACTGATACTGTTCCTGGCACTGTTGAGGTAGTTGAGCCTGAAGCT |
| 9 | GGAACTGTCACTACTACTGTCTATTCTGGTACTCAAGAGTACACTACCACACTTGCTACAGCCAGTGGTACTGTTTCAGGCACTGTTGAAGTAGTGGATACAGCAGCT |
| 10 | GGAACTGTTACTACTACAATTTACTCTGGAACTACACCATTCAACACAACACTTGCTTCCGCAACTGATACTGTTCCTGGCACTGTTGAGGTAGTTGAGCCTGAAGCT |
| 11 | GGAACTGTCACTACTACTGTATATTCTGGCACTCAAGAGTACACCACAACGCTTGCTACAGCTAGTGGTACTGTTTCAGGCACTGTTGAAGTAGTGGATACAGCAGCT |
| 12 | GGAACTGTTACTACTACAATTTACTCTGGAACTACACCATTCAACACAACACTTGCTTCCGCAACTGATACTGTTCCTGGCACTGTTGAGGTAGTTGAGCCTGAAGCT |
| 13 | GGAACTGTCACTACTACTGTCTATTCTGGTACTCAAGAGTACACCACAACGCTTGCTACAGCTAGTGGTACTGTTTCAGGCACTGTTGAAGTAGTGGATACAGCAGCT |
| 14 | GGAACTGTTACTACTACAATTTACTCTGGAACTACACCATTTAACACAACGCTTGCTTCCGCAACTGATACTGTTCCTGGCACTGTTGAGGTAGTTGAGCCTGAAGCT |
| 15 | GGAACTGTCACTACTACTGTATATTCTGGCACTCAAGAGTACACTACCACACTTGCTACAGCCAGTGGTACTGTTTCAGGTACCGTTGAAGTAATTGAACCTGCTGCA |
| 16 | GGAACCGTCACTACTACTGTATATTCTGGCACTCAAGAGTACACCACAACGCTTGCTACAGCCAGTGGTACTGTTTCAGGTACCGTTGAAGTAATTGAACCTGCTGCA |
| 17 | GGAACCGTTACTAGTACTGTCTATTCTGGTACTCAAGAGTACACCACAACGCTTGCTACAGCCAGTGGTACTGTTTCAGGCACCGTTGAAGTAATTGAACCCGCTGCA |
| 18 | GGAACCGTTACTACTACTGTTTACTCTGGTACTCAAGAGTACACTACCACACTTGCTACAGCCAGTGGTACTGTTTCAGGCACCGTTGAAGTAATTGAACCCGCTGCA |
| 19 | GGAACCGTTACTACTACTGTCTATTCTGGTAGCGAGGTATATACTACAACTCTCGCTTCTGCTTCCGAATCTGTCCCCGGAACTGTTGAAGTAGTAGACCCAGAAGCT |
| 20 | GGAAGTGTCACCACTACTATCTATTCTGGTAGCGTAGAGTACACTACTGTTCTTGCTGATGCAAGTGGATCTGTTACTGGGACTGTTGAAGTTGTTGAACCGGCTGCA |
| 21 | GGAACTGTCACCACTACTGTTTACTCTGGTAGCGAGGTATATACTACAACTCTTGCTTCTGCTTCCGAATCTGTCCCCGGAACTGTTGAAGTAGTAGACCCAGAAGCT |
| 22 | GGAAGTGTCACCACTACTATCTATTCTGGTAGCGTAGAGTACACTACTGTTCTTGCTGATGCAAGTGGATCTGTTACTGGAACTGTTGAAGTAGTTGAACCTGCTGCT |
| 23 | GGAACTGTTACCGGTACCCTCACTTCCGGTTCTCAGTTTTTTACTACTACTATCGCTCAAGCTAGTGGTTCCGTATCCGGCAACGTGGAGGTAATTGAGCCTTCTGGT |
| 24 | TCAACTGTTACTTCAACTATCTATAGTGGTAGTGAATCTTTCACTACGACTCTTGCTGTTGGAAGTGGAACTATTCCAGGTACTGTGGAGGTAATTCTCCCCGCTCCT |

**SPCC188.09C- *pfl4* Nucleotide sequence**

| | Sequence |
|---|---|
| 1 | AGCACAATATACACCACAATCAACAGCGGTACTGTAGCTTCCACATATACTGTCACACTGGCAGATGGCGACGTT------GTGGTCAAAGACATAGAGCCCACTGCA |
| 2 | GGAACGGTGACTACCACGATCACTTCAGGTAGTCACGAGTTCACTACCACTCTTGCTGAAGCCAGTGGTACTGTCCCTGGAACGGTTGAGGTAGTTGAGCCTCTAGCT |
| 3 | GGAACTGTTACTACTACTATTCATTCCGGTAGTGTAGAATACAACACCACTCTTGCTACCGCAAGTGGAACTGTTCCTGGTACCGTTGAGGTTGTTGAACCCGCTGTT |
| 4 | GGAACTGTTACTACTACCATATACTCAGGCAGTGAAGAGTTTACAACTACTTTGGCATCTGCCTCTGGTTCAATTTCAGGTACCGTTGAAGTAATTGAACCTACAGCT |

| | |
|---|---|
| 5 | GGCACAGTAACTACCACTATTTACTCTGGCGACCAAGAGTATACCACTATCCTTGCTGAAGCCAGTGGAACTGTTCCAGGTACTGTTGAAGTGATTGAACCTGCTGTT |
| 6 | GGAACCGTTACGACCACTACTTACTCAGGTAGTGTAGAGTATACAACTACTTTGGTACCTGCCTCTGGTTCTGTCTCCGGAACCGTTGAGGTCGTTGAGCCTGCGGTT |
| 7 | GGAACTGTGACGACGACATTGCAGTCTGGGTCTCAAGCCTTTACAACCACTGTG---CCCGCCTCCGGTAGTGTATCGGGAACTGTTGAGGTAGTTCAGCCAACTGGT |
| 8 | GGCACTGTTACTAATACAGTATATGAGGGATCTCAGACCATTACTTCGACTCTTGCAACAGCTAGTGGTACTGTTCCAGGTACTGTTGAAGTTATTCTTCCTGGGCCT |
| 9 | AGCACAATCTAT------------TCGGGAACTGTTGCTACCACTATTACATATGATGTTTCTAGTACGCCTGCGTCTACTGTTGTTGTT---ATC---CCTACTGCT |

**Supplemental Figure S6.** An alignment of the nucleotide sequences of *pfl*10 and *pfl*4 shows their homolog

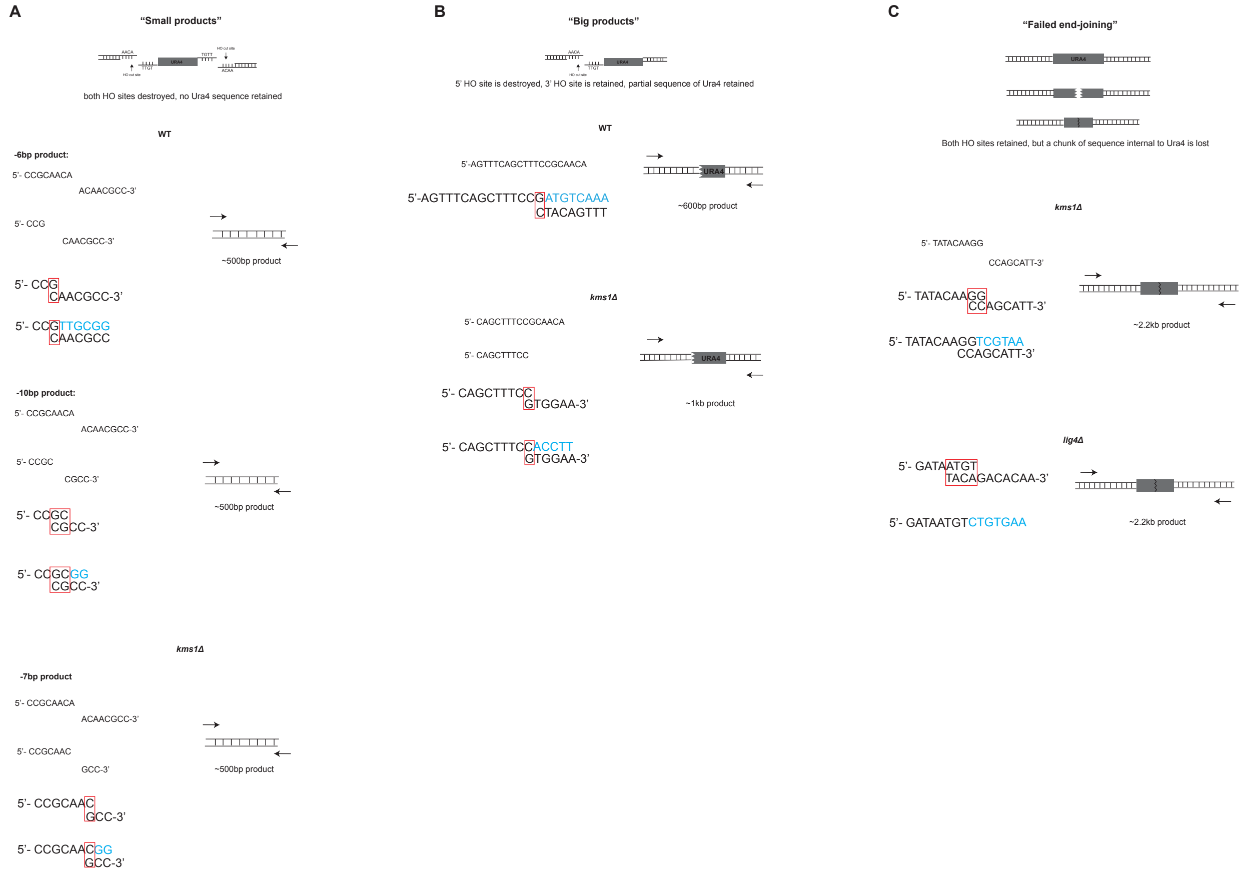

**Supplemental Figure S7. Recombinants can maintain portions of *ura4* by repairing with microhomologies within the *ura4* coding sequence.** A) "Small products" are individual FOA+ products where both HO sites are destroyed. These products are outlined in Figure 7D. Two of the WT products (the -6bp product and the -10bp product) are possible via rejoining of a 1-2bp microhomology. One *kms1Δ* product (the -7bp product) is also a possible via a 1bp microhomology. B) "Large products" are products where the HO site 5' to *ura4* is destroyed, but the HO site 3' to *ura4* is retained. These are possible via repair by a 1bp microhomology within the *ura4* coding sequence. Interestingly, the product size in WT is similar to the dual cutters, making it difficult to distinguish between them in an ensemble of FOA+ colonies, as in Fig. 7C. In *kms1Δ* cells we observe many examples that retain more of the sequence, which explains the variety of product sizes seen in Fig. 7C. C) "Failed cNHEJ" products have both HO sites intact, are not observed in WT cells, and arise in cells lacking *lig4*, suggesting that they are unable to complete cNHEJ. These examples lost ~300bp of sequence internal to *ura4* (resulting in a ~2.2kb product), and the junction is a result of rejoining of microhomologies. Similar products are observed in *kms1Δ* cells where repair appears to occur via a 2bp microhomology.

**Supplemental Table S1:** Rates of recombination as mean ± standard deviation.

| **Intragenic recombination** | |
|---|---|
| **Genotype** | **Rate of recombination** |
| *pfl3:3'kan-ura4+* | $4.39 \times 10^{-6} \pm 9.88 \times 10^{-7}$ |
| *pfl10:ura4+-5'kan* | $1.41 \times 10^{-5} \pm 3.55 \times 10^{-6}$ |
| *pfl5:ura4+-5'kan* | $1.53 \times 10^{-5} \pm 5.35 \times 10^{-6}$ |
| *map4 200-3:ura4+-5'kan* | $1.59 \times 10^{-6} \pm 8.38 \times 10^{-7}$ |
| *mam3 201-1:3'kan-ura4+* | $5.25 \times 10^{-6} \pm 4.22 \times 10^{-6}$ |
| *pfl5:3'kan-ura4+ lig4Δ::kanMX6* | $7.83 \times 10^{-6} \pm 8.34 \times 10^{-7}$ |
| *pfl5:ura4+-5'kan exo1Δ::kanMX6* | $1.14 \times 10^{-5} \pm 1.56 \times 10^{-6}$ |
| *pfl5:ura4+-5'kan rad52Δ::kanMX6* | $6.70 \times 10^{-5} \pm 4.88 \times 10^{-5}$ |
| *pfl5:ura4+-5'kan rad51Δ::hygMX6* | $8.17 \times 10^{-4} \pm 1.19 \times 10^{-4}$ |
| *pfl5:ura4+-5'kan rad55Δ::hygMX6* | $1.22 \times 10^{-4} \pm 1.68 \times 10^{-5}$ |
| *pfl5:ura4+-5'kan swi10Δ::kanMX6* | $4.39 \times 10^{-5} \pm 2.2 \times 10^{-5}$ |
| *pfl5:3'kan-ura4+ cbp1Δ::kanMX6* | $5.06 \times 10^{-5} \pm 1.68 \times 10^{-6}$ |
| *pfl3:3'kan-ura4+ cbp1Δ::kanMX6* | $1.14 \times 10^{-5} \pm 1.32 \times 10^{-6}$ |
| *pfl10:3'kan-ura4+ cbp1Δ::kanMX6* | $1.61 \times 10^{-5} \pm 3.19 \times 10^{-6}$ |
| *pfl10:ura4+-5'kan tf2-3Δ::kanMX6* | $2.16 \times 10^{-5} \pm 1.03 \times 10^{-5}$ |
| *pfl5:3'kan-ura4+ kms1Δ::natMX6* | $2.39 \times 10^{-5} \pm 9.35 \times 10^{-6}$ |
| *pfl5:3'kan-ura4+ kms1Δ::natMX6 cbp1Δ::kanMX6* | $4.73 \times 10^{-5} \pm 1.82 \times 10^{-5}$ |
| *natMX-nmt41:pfl5:ura4+-5'kan (uninduced)* | $3.01 \times 10^{-5} \pm 1.76 \times 10^{-6}$ |
| *natMX-nmt41:pfl5:ura4+-5'kan (induced)* | $2.65 \times 10^{-5} \pm 4.62 \times 10^{-6}$ |
| **Intergenic recombination** | |
| **Genotype** | **Rate of recombination** |
| *pfl10:ura4+-5'kan pfl4:3'kan-ura4+* | $2.09 \times 10^{-8} \pm 5.61 \times 10^{-9}$ |
| *pfl10:ura4-5'kan pfl4:3'kan-ura4 kms1Δ::hygMX6* | $6.24 \times 10^{-8} \pm 1.45 \times 10^{-8}$ |
| *pfl10:ura4-5'kan pfl4:3'kan-ura4 cbp1Δ::hygMX6* | $6.03 \times 10^{-8} \pm 3.98 \times 10^{-8}$ |
| **Repair of incompatible ends and loss of functional *ura4*** | |
| **Genotype** | **Rate of repair** |
| *mmf1:HOcs-ura4+-HOcs* | $1.27 \times 10^{-6} \pm 3.27 \times 10^{-6}$ |
| *mmf1:HOcs-ura4+-HOcs lig4Δ::natMX6* | $6.78 \times 10^{-7} \pm 1.37 \times 10^{-7}$ |
| *mmf1:HOcs-ura4+-HOcs pku70Δ::natMX6* | $1.36 \times 10^{-6} \pm 8.10 \times 10^{-6}$ |
| *mmf1:HOcs-ura4+-HOcs kms1Δ::natMX6* | $3.84 \times 10^{-6} \pm 4.24 \times 10^{-6}$ |

**Supplemental Table S2:** Strains used in this study.

| Strain | Genotype |
|---|---|
| MKSP1019 | *h- ade6? ura4-D18 leu? sad1-mCherry:kanMX6 rad52-GFP:kanMX6* |
| MKSP1184 | *h- ade? ura4-D18 leu? sad1-mCherry:kanMX6 rad552-GFP:kanMX6 kms1Δ::hygMX6* |
| MKSP1615 | *h- ade6-M216 ura4-D18 leu1-32 pfl10:ura4+-5'kanMX6 pfl4:3'kanMX6-ura4+* |
| MKSP1616 | *h+ ade6-M216 ura4-D18 leu1-32 pfl10:ura4+-5'kanMX6 pfl4:3'kanMX6-ura4+ kms1Δ::hygMX6* |
| MKSP1652 | *h+ ade6-M216 ura4-D18 leu1-32 pfl5:ura4+-5'kan* |
| MKSP1655 | *h? ade6-M216 ura4-D18 leu1-32 pfl5:3'kan-ura4+ cbp1Δ::kanMX6* |
| MKSP1658 | *h+ ade6-M216 ura4-D18 leu1-32 natMX-pnmt41:pfl5:ura4+-5'kan* |
| MKSP1661 | *h- pfl5:lacO-ura4+ lacI-GFP cut11-mCherry:natMX6* |
| MKSP1663 | *h? pfl10:lacO-ura4+ lacI-GFP cut11-mCherry:natMX6* |
| MKSP1668 | *h? pfl10:lacO-ura4+ lacI-GFP cut11-mCherry:natMX6 cbp1Δ::hygMX6* |
| MKSP1713 | *h? ade6-M216 ura4-D18 leu1-32 pfl10:ura4+-5'kan pfl4:3'kan-ura4 cbp1Δ::hygMX6* |
| MKSP1714 | *h- ade6-M216 ura4-D18 leu1-32 pfl3:3'kan-ura4+* |
| MKSP1838 | *h- mam3:lacO-ura4+ lacI-GFP cut11-mCherry:natMX6* |
| MKSP1839 | *h+ ade6-M216 ura4-D18 leu1-32 pfl10:ura4+-5'kan tf2-3Δ::kanMX6* |
| MKSP1841 | *h? ade6-M216 ura4-D18 leu1-32 pfl5:3'kan-ura4+ lig4Δ::kanMX6* |
| MKSP1855 | *h? ade6-M216 ura4-D18 leu1-32 pfl5:ura4+-5'kan exo1Δ::kanMX6* |
| MKSP1858 | *h? ade6-M216 ura4-D18 leu1-32 pfl5:ura4+-5'kan rad52Δ::kanMX6* |
| MKSP1880 | *h- ade6-M216 ura4-D18 leu1-32 pfl3:3'kan-ura4+ cbp1Δ::kanMX6* |
| MKSP1882 | *h- ade6-M216 ura4-D18 leu1-32 pfl10:3'kan-ura4+* |
| MKSP1884 | *h? ade6-M216 ura4-D18 leu1-32 pfl5:ura4+-5'kan swi10Δ::kanMX6* |
| MKSP1888 | *h? ade6-M216 ura4-D18 leu1-32 pfl5:ura4+-5'kan rad51Δ::hygMX6* |
| MKSP1895 | *h? ade6-M216 ura4-D18 leu1-32 pfl10:3'kan-ura4+ cbp1Δ::kanMX6* |
| MKSP1938 | *h- map4:lacO-ura4+ lacI-GFP cut11-mCherry:natMX6* |
| MKSP1944 | *h+ ade6-M216 ura4-D18 leu1-32 pfl5:ura4+-5'kan rad55Δ::hygMX6* |
| MKSP1966 | *h+ ade6-M216 ura4-D18 leu1-32 map4 200-3:ura4+-5'kan* |
| MKSP1974 | *h- ade6-M216 ura4-D18 leu1-32 mam3 201-1:3'kan-ura4+* |
| MKSP1983 | *h? pfl5:lacO-ura4+ lacI-GFP cut11-mCherry:natMX6 cbp1Δ::kanMX6* |
| MKSP2018 | *h- leu1-32 ura4-D18 ade6? lys1? his7::GFP-LacI-NLS-his7 cut3-477::lacO-cut3+ cut11-mCherry::natR* |
| MKSP2310 | *h- pfl3:lacO-ura4+ lacI-GFP cut11-mCherry:natMX6* |
| MKSP2354 | *h? pfl3:lacO-ura4+ lacI-GFP cut11-mCherry:natMX6 cbp1Δ::kanMX6* |
| MKSP3795 | *h- ade6? ura4-D18 leu? sad1-mCherry:kanMX6 rad52-GFP:kanMX6 rad51Δ::HygMX6* |
| MKSP3796 | *h- ade6? ura4-D18 leu? sad1-mCherry:kanMX6 rad52-GFP:kanMX6 rad51Δ::HygMX6 kms1Δ:: HygR* |
| MKSP4299 | *h- ade6+ leu1-32 ura4-D18 mmf1-HOcs-ura4+-HOcs* |
| MKSP4305 | *h- ade6+ leu1-32 ura4-D18 mmf1-HOcs-ura4+-HOcs kms1Δ::natMX6* |
| MKSP4313 | *h- ade6+ leu1-32 ura4-D18 mmf1-HOcs-ura4+-HOcs lig4Δ::natMX6* |

| | |
|---|---|
| MKSP4362 | *h- ade6+ leu1-32 ura4-D18 mmf1-HOcs-ura4+-HOcs pku70Δ::natMX6* |
| MKSP4434 | *h? ade6-M216 ura4-D18 leu1-32 pfl5:3'kan-ura4+ kms1Δ::natMX6* |
| MKSP4435 | *h? ade6-M216 ura4-D18 leu1-32 pfl5:3'kan-ura4+ kms1Δ::natMX6 cbp1Δ::kanMX6* |

Note: precise insertion sites of the *ura4* and *kanR* cassette fragments are uncertain due to the repetitive nature of the *pfl* genes. For approximate locations please see Supplemental Table S3.

**Supplemental Table S3:** Insertion sites for LacO and *ura4*.

| Genetic locus | LacO insertion location |
|---|---|
| *cut3* | chrII:1006688-1002263 |
| *pfl5* | chrII; 4403547-4403553 |
| *pfl10* | chrI; 2959337-2961046 |
| *pfl3* | chrII; 665084-667124 |
| *mam3* | chrI; 1837071- 1838471 |
| *map4* | chrII; 2450913-2452027 |

| Genetic locus | Approximate *ura4* insertion location |
|---|---|
| *pfl5* | chrII; 4421537-4424856 |
| *pfl10* | chrI; 2959337-2961046 |
| *pfl3* | chrII; 665084-667124 |
| *mam3* | chrI; 1837071- 1838471 |
| *map4* | chrII; 2450913-2452027 |

The position of *ura4* insertion sites within the repeats of the *pfl* genes have been approximately determined based on PCR analysis.

**Supplemental Table S4:** Primers used in this study.

| Primer name | Sequence |
|---|---|
| Mam3_For | 5'-TGGTTCCGTTGGATTTACC-3' |
| Mam3_Rev | 5'-CAGAATGGTGAGGTGTTCC-3' |
| Map4_For | 5'- GTGTCTTCATCTCATCCGTATC-3' |
| Map4_Rev | 5'- TATCGTCCTGGTCCAACCT-3' |
| PFL5_For | 5'- CCACACTCAACAATAAGCACTC-3' |
| PFL5_Rev | 5'- GCGGGAACAGTAGTAGTAAACC-3' |
| PFL3_For | 5'- TAGTGGAACAGTCAGCGGT -3' |
| PFL3_Rev | 5'- CAGCCAGGGAATAAAGAAGG -3' |
| PFL10For | 5'- ACTCTTGACTCGGCAAATG-3' |
| PFL10Rev | 5'- AACAGTTGAACCAGAAGGCT-3' |
| PFL4For | 5'- TGTGGTCAAAGACATAGAGCC-3' |
| PFL4Rev | 5'- GGATAACAACAACAGTAGACGC-3' |
| Act1_For1_qP | 5'- CCAAATCCAACCGTGAGAAGA -3' |
| Act1_Rev1_qP | 5'- GTACGACCAGAGGCATACAAAG -3' |
| Act1_For2_qP | 5'- ACGTCGCTTTGGACTTTGA -3' |
| Act1_Rev2_qP | 5'- CGCTCGTTTCCGATAGTGATAA -3' |
| PFL3_For1_qP | 5'- CAACCTCTTCTCAGCGACATAA -3' |
| PFL3_Rev1_qP | 5'- ACATCATCGGTCCCTGTAGTA -3' |
| PFL3_For2_qP | 5'- CTACAGGGACCGATGATGTTG -3' |
| PFL3_Rev2_qP | 5'- AGTTGTCGTCGGATTTGTAGG -3' |
| PFL5_For1_qP | 5'- CGTCTGGTGCTGTCATAGTT -3' |
| PFL5_Rev1_qP | 5'- GGATAGTCGAGGTAAACGGAATAG -3' |
| PFL5_For2_qP | 5'- GCTGGACTTCAACGGTAACT -3' |
| PFL5_Rev2_qP | 5'- ACACCAGCACTACACCAATC -3' |
| PFL10_For1_qP | 5'- CTCAAGCTAGTGGTTCCGTATC -3' |
| PFL10_Rev1_qP | 5'- CAGCAAGAGTCGTAGTGAAAGA -3' |
| PFL10_For2_qP | 5'- TTGCTAATGGAGGTGGAAAGG -3' |
| PFL10_Rev2_qP | 5'- TGAGTGCAAGTAGCTGTGTAAG -3' |
| Mmf1_L1 | 5'- GCATTTTGTCGATGATGAACGTTG-3' |
| Mmf1_L1Seq | 5'- CGGTTGGATACATCTCTGTCATC-3' |
| Mmf1_L6 | 5'- GGCGCTTTTCATTTTCGTGC-3' |
| Aly3F | 5'- CGTGTTATTAGGACTATTCCTC-3' |
| Aly3R | 5'- CCTACGTAGCAGTAAGAATACAGC-3' |
| Ura4_ins_AscI | 5'-CAGTCAGGCGCGCCAGCTTAGCTACAAATCCCACTG-3' |
| Ura4_ins_PmeI | 5'- CTGACAGTTTAAACGGTCGACGGTATCGATAAGC-3' |
| Ura_int_chk | 5'- GCCAGCTGAAGCTTCGTAC-3' |
| HOcs_ins_AscI_F | 5'- TTGGCGCGCCAGTTTCAGCTTTCCGCAACAGTATAATTTTATAA AGGCGCGCCTT-3' |
| HO_ins_AscI_R | 5'- AAGGCGCGCCTTTATAAAATTATACTGTTGCGGAAAGCTGAA ACTGGCGCGCCAA-3' |

**Supplemental Table S5:** Plasmids used in this study.

| Plasmid name | Method of generation | Description |
|---|---|---|
| pSR10_ura4_5.6kb | Previous work (Leland & King, 2014) | Integrated nearby the *pfl* loci and *cut3* to visualize their localization within then nucleus |
| pFA6a_HOcs_Ura | Plasmid was cut at AscI and PmeI to release the Hyg-resistance gene fragment. An insert with the *S. pombe ura4* gene was created by amplifying *ura4* from a pFA6a-Ura4. | Used plasmid from a previous study (Leland et al., 2018) which contained a modified MX6-based hygromycin-resistance cassette containing the HO cut site to switch the marker from hygromycin to *ura4*. |
| pFA6a_HOcs_Ura_HOcs | Added another HO endonuclease cut site at the AscI site of pFA6a_HOcs_Ura in the inverted orientation. | Integrated into the *mmf1* locus of *S. pombe* for the salvage DNA repair assay. |
